# A Surfactant Cocktail Overcomes Air-Water Interface Artifacts in Single-Particle CryoEM

**DOI:** 10.64898/2026.03.17.712260

**Authors:** Suzanne E. Enos, Brian D. Cook, Hamidreza Rahmani, Sarah M. Narehood, Yizhou Li, Inga C. Kuschnerus, Trevor H. Redford, Peter Dukakis, Daniel Ji, Maxwell J. Bachochin, Danielle A. Grotjahn, Mark A. Herzik

## Abstract

Single-particle cryogenic electron microscopy (cryoEM) is a widely used technique for structure determination of biomacromolecules to near-atomic resolution. Random distributions of these molecules in vitrified ice are necessary to accumulate enough two-dimensional views to generate a complete three-dimensional (3-D) reconstruction. However, interactions between the sample and the air-water interface (AWI) that occur during vitrification often bias the views of the sample, a phenomenon termed preferred orientation, limiting our ability to obtain 3-D reconstructions. Surfactants are often used as sample additives to prevent AWI-induced deterioration, but no general strategy exists for surfactant choice, requiring laborious screening for each sample. To circumvent these issues, we developed SurfACT, a cocktail of diverse surfactants with distinct physicochemical properties that limits AWI-dependent sample denaturation and orientation bias, while mitigating individual surfactant-specific drawbacks. Here we demonstrate SurfACT’s effectiveness with four proteins plagued by AWI-induced issues, including two species of hemagglutinin (HA), molybdenum-iron protein (MoFeP) from the nitrogenase enzyme, and aldolase. All four samples show drastically improved viewing distribution and map completeness when SurfACT is applied. Cryogenic electron tomography demonstrates that SurfACT redistributes particles from the AWI into the bulk ice, driving signal recovery and inhibiting denaturation. This versatile sample additive minimizes sample-specific screening and expands the capabilities and range of suitable samples for cryoEM.

## Introduction

In single-particle cryogenic electron microscopy (cryoEM), obtaining two-dimensional (2-D) projection images (i.e., micrographs) of randomly distributed biomacromolecules (i.e., particles) in a thin layer of vitreous ice is necessary to accumulate sufficient distinct 2-D views to generate a complete three-dimensional (3-D) reconstruction.^1,2^ Critically, the process of vitrifying cryoEM samples to obtain enough 2-D views to sufficiently sample 3-D space has been a major hurdle within the single-particle cryoEM (SPA) field for decades.^3–7^ While the major underlying molecular and physical mechanism leading to the loss of these views, a phenomenon broadly referred to as preferred orientation, remains incompletely understood, the outcome of biased 2-D projections of the sample is well documented — incomplete sampling of Fourier space and consequent loss of density in the final volume, manifesting as poor local map quality or entire missing domains.^8–13^

Numerous approaches have been undertaken to address the preferred orientation problem in cryoEM, each tailored to reconcile different stages of the sample preparation workflow hypothesized to deteriorate sample quality, all with mixed successes and noted limitations.^14^ The prevailing hypothesis is that particles interact with the air-water interface (AWI) during the vitrification process, leading to the destabilization of (sub-)domains and/or the entire protein, resulting in biased and persistent interactions with the AWI and preferential orientation of these molecules upon vitrification.^1,10,15^ The underlying idea is that if interactions with the AWI are eliminated, or the AWI is removed entirely, these problems can be ameliorated, resulting in improved view sampling, more complete 3-D reconstructions, and higher-quality EM maps.

Solid support layers like graphene, graphene oxide, or amorphous carbon support films overlaid on the EM grid surface have been demonstrated to limit the effects of one of the AWIs, but preparation of these support films often presents batch-to-batch variations, and great care must be taken to ensure that the sample is not overwicked unknowingly, resulting in sample dehydration and denaturation.^16–24^ The development of rapid automated plunge-freezing, including the Spotiton and SPT Labtech chameleon, have been demonstrated to improve the quality of some samples, speculated as the ability of the instrumentation to ‘outrun’ AWI association, but these instruments tend to exhibit highly sample-dependent improvements.^25–34^ For some pathologically preferred specimens, tilted data collection, which involves physically adjusting the angle of the grid within the microscope (e.g., 20-40°) during data collection, can access more views, but the thicker ice cross-section results in increased beam-induced motion and decreased signal-to-noise that limits the attainable resolution.^35,36^ Machine learning algorithms offer a computational solution that attempts to “recover” data from missing views through inference but can suffer from hallucinations and misleading information unless proper validation is pursued.^37–40^

Inclusion of surfactants has become a common strategy to reduce deleterious particle interactions with the AWI by forming a physical barrier with the AWI to avoid sample interactions.^41^ These additives, ranging from non-ionic detergents to amphipathic polymers, have been demonstrated to improve particle orientation distributions, reduce denaturation at the AWI, and, in some cases, increase overall particle yields (**Figure 1**).^41–45^ Although single additives can work for specific samples, particularly when optimized carefully for concentration and buffer compatibility, success often requires extensive sample-dependent screening to maintain folded protein and intact complexes.^41^

**Figure 1:**
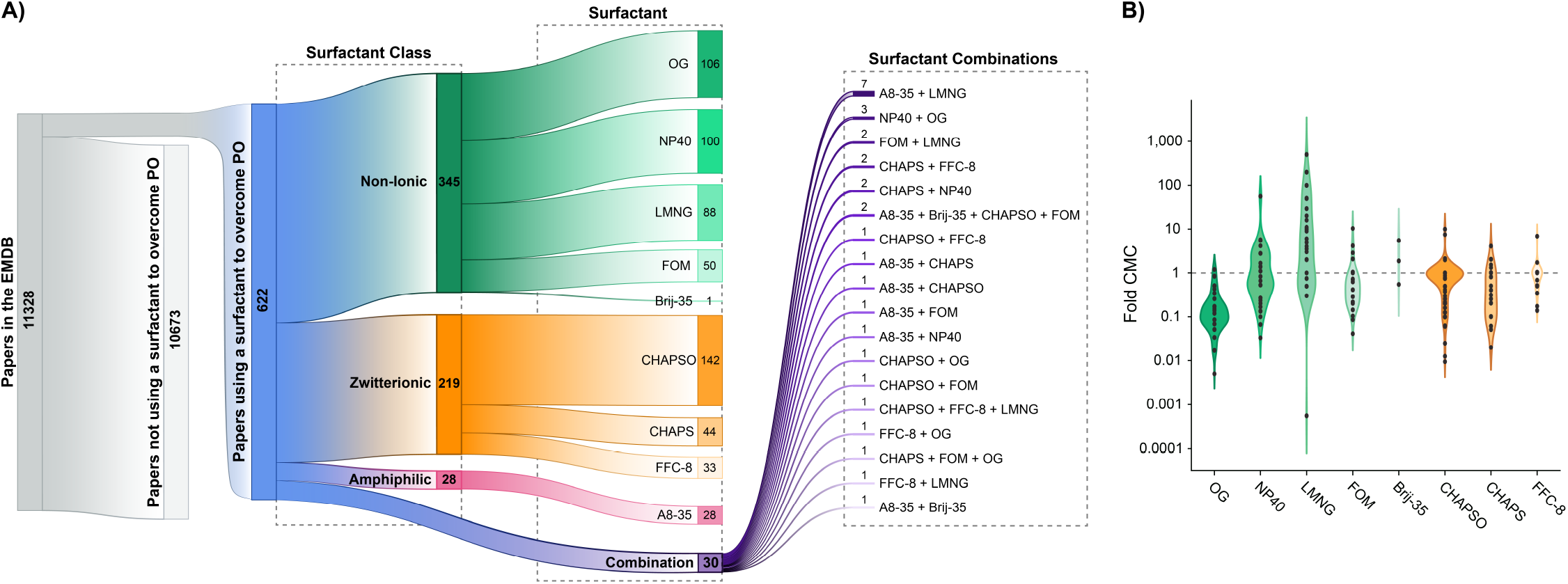
Systematic Survey of Surfactant Additive Use in Deposited CryoEM Structures. **A**) Sankey plot showing documented surfactant additives for preferred orientation (PO) in publications referenced in the EMDB. Nodes separate publications using surfactant(s) to overcome preferred orientation (*left*), surfactant class by chemical properties (*middle*), and surfactant identity (*right*). **B**) Violin plot showing concentrations of all single-additive use cases relative to the critical micelle concentration (CMC) for each respective surfactant: octyl glucoside (OG), nonidet P-40 (NP40), lauryl maltose neopentyl glycol (LMNG), fluorinated octyl maltoside (FOM), Brij-35, 3-[(3-cholamidopropyl)-dimethylammonio]-2-hydroxy-1-propanesulfonate (CHAPSO), 3-[(3-cholamidopropyl)-dimethylammonio]-1-propane sulfonate (CHAPS), and fluorinated fos-choline-8 (FFC-8). Amphipol 8-35 (A8-35) was excluded from this analysis since it does not possess a traditional CMC.

In practice, the effects of individual surfactants are often stochastic and unpredictable, with many complexes exhibiting destabilization, aggregation, or conformational perturbation at higher surfactant concentrations, or, in some cases, surfactants directly binding to the surface of the protein under evaluation.^14,44,46^ These limitations complicate analyses that restrict broad applicability and make single-additive strategies powerful but variable, especially for fragile or orientation-prone complexes, while imposing a high experimental burden that scales poorly when screening across multiple targets or preparation modalities.

A useful conceptual precedent for addressing the issues incurred by single-additive screening comes from X-ray crystallography, where cryoprotectant selection has long faced analogous challenges.^47^ Early cryo-cooling studies showed that using binary or ternary mixtures of cryoprotectants (e.g., glycerol combined with one or more various additives) could achieve vitrification at lower total solute concentrations, reduce ice formation, and minimize osmotic or mechanical stress compared with using any single agent at high concentration.^48^ In principle, these mixtures work synergistically with each component contributing distinct physicochemical effects, such as lowering the freezing point, suppressing ice nucleation, reducing local heterogeneity, and preserving crystal integrity.^47,48^

We sought to apply this principle to cryoEM sample preparation: rather than forcing a single additive to perform multiple incompatible roles, a rationally designed, low-concentration mixture of surfactants could achieve robust and uniform AWI coverage while minimizing deleterious effects on the biomacromolecule of interest. By leveraging complementary surface chemistries, such a cocktail could provide more predictable and generalizable mitigation of AWI interactions, reduce the need for extensive condition screening, and expand the range of complexes amenable to high-resolution analysis.

Here, we present a rationally designed surfactant cock-tail termed SurfACT (**SURF**actants for **A**ir-water interface **C**on**T**rol), based on a comprehensive analysis of surfactant use in the Electron Microscopy Data Bank (EMDB) and inclusion of diverse surfactant chemistries. We demonstrate SurfACT’s effectiveness with four proteins plagued by preferred orientation and/or AWI-dependent loss of sample integrity: two diverse influenza glycoprotein hemagglutinins (HA), molybdenum-iron protein (MoFeP) from *Azotobacter vinelandii* nitrogenase, and rabbit muscle aldolase. In all four cases, SurfACT improved viewing distribution, enhanced volume completeness, and reduced or eliminated sample destabilization. We further show that SurfACT requires minimal sample-specific optimization, decreases reliance on enforced symmetry in data processing to generate complete 3-D reconstructions, and increases high-quality particle yields during data collection, making it a useful general strategy for cryoEM sample preparation.

## Results

### A Systematic Analysis of Surfactant Use in CryoEM Sample Preparation

While individual reports have demonstrated that specific surfactants can improve particle distribution and/or sample integrity for particular specimens, the broader landscape of surfactant use across cryoEM studies remains poorly defined.^10^ Furthermore, inconsistent documentation in the EMDB complicates these evaluations. To our knowledge, no systematic survey of deposited cryoEM structures has examined how frequently surfactants are employed as a strategy to mitigate AWI-induced artifacts, or how these additives are distributed across surfactant classes or use cases.

To assess how surfactant additives have been employed in practice, we surveyed entries from the EMDB for deposited structures that explicitly report the use of detergents, surfactants, or amphiphilic polymers in the cryoEM sample preparation steps of their associated publications (**Supplementary Table 1**). Using EMDB metadata in combination with manual curation of associated “Methods” sections from publications, we identified 622 entries that listed one or more additives introduced specifically to address preferred orientation or AWI-related issues. These additives include non-ionic detergents (e.g., octyl glucoside (OG), nonidet P-40/NP40, lauryl maltose neopentyl glycol (LMNG), fluorinated octyl maltoside (FOM)), zwitterionic detergents (e.g., 3-[(3-cholamidopropyl)-dimethylammonio]-1-propane sulfonate (CHAPS), 3-[(3-cholamidopropyl)-dimethylammonio]-2-hydroxy-1-propanesulfonate (CHAPSO), fluorinated fos-choline-8 (FFC-8)), and amphiphilic polymers (e.g., A8-35), among others, which we categorized based on additive class (**Figure 1A**).

Several clear trends emerged from this analysis. First, the overwhelming majority of the studies (*>*95% of the 622 publications) employed single additives to mitigate AWI problems, with most manuscripts (55%) reporting the use of non-ionic detergents. Zwitterionic detergents were also commonly employed but less frequently than non-ionic detergents, while amphiphilic polymers are comparatively rare. These cases demonstrate that single additive success can be achieved, but a wide variety of options must be screened due to little consensus in the literature on best practices. Second, less than 5% of the survey entries (30 entries in total) report the use of more than one surfactant additive, with 12 examples reporting one-off mixtures not used in any other studies. Unfortunately, the fact that unsuccessful use of either individual surfactants or combinations are almost never reported limits our ability to fully evaluate the relative benefits of one surfactant class over another. Collectively, our analysis of the EMDB underscores the lack of a single universally effective surfactant or surfactant mixture and highlights the need for a rationally designed surfactant combination that balances interfacial behaviors while minimizing destabilizing effects (**Figure 1A**).

Next, we examined the concentration at which single-use surfactants were employed relative to their critical micelle concentration (CMC), since micelles can complicate particle picking and contribute to electron scattering to reduce signal-to-noise, and surfactant concentrations near or above the CMC increase the likelihood of disrupting protein fold or complex interactions.^49^ In general, we observed surfactants used to combat AWI issues are supplied between 0.1-fold and 10-fold the CMC, with LMNG skewing towards higher concentrations, likely due to its very low CMC, and OG use favoring slightly lower concentrations (**Figure 1B**). This wide range of reported surfactant concentrations reveals another level of inconsistency that must be evaluated when facing preferred orientation and AWI-induced artifacts. In formulating a surfactant cocktail, we sought to minimize deleterious destabilizing effects of the individual components by keeping concentrations as low as possible while remaining effective. From our survey, we found that for most surfactants the lowest concentration with documented success was 0.1-fold CMC, so we elected to use this as the starting point for our surfactant cocktail formulation.

### SurfACT Formulation, Application Guidelines, and Considerations for Use

Trends from the single-surfactant use survey directly motivated our development of a mixed-surfactant cocktail, designed to harness the beneficial aspects of individual agents while counterbalancing their detrimental and stochastic effects. To assemble our sample additive cocktail (SurfACT) we selected four mild, protein-stabilizing surfactants — avoiding membrane extraction capabilities (OG, NP40, LMNG) — with diverse physical and chemical properties that have each been demonstrated to reduce preferred orientation and/or AWI-induced sample degradation in cryoEM: FOM, a small fluorinated non-ionic detergent; Brij-35, a large non-ionic detergent; CHAPSO, a zwitterionic detergent; and A8-35,^50^ an amphiphilic polymer (**Figure S1**). These components’ varied physical and chemical properties, and the use of each at 0.1-fold CMC (in the case of A8-35, 0.1-fold average protein-solubilizing concentration), is expected to prevent micelle formation (which can lead to false particle picks during cryoEM data processing)^51–54^ while providing diverse interactions with the sample, limiting background scattering, and decreasing grid freezing pathologies. When used individually, these surfactants would require sample-dependent optimization to ensure sample integrity, but our combinatorial additive approach strives to circumvent such issues and facilitate broad applicability to a wide variety of proteins affected by preferred orientation.

To avoid exposing samples to surfactant micelles or phase partitions, we pre-dilute SurfACT into sample buffer prior to use, then mix the sample and SurfACT 1:1 (v/v) immediately prior to grid preparation (see **Methods**). This approach minimizes the formation of local surfactant-rich microenvironments which can transiently expose sample to surfactant concentrations far exceeding their effective working concentration, destabilizing weak protein-protein interactions, or perturbing conformational equilibria. By carefully mixing 1:1, we ensure the sample encounters a spatially uniform surfactant environment, improving reproducibility and reducing the likelihood of surfactant-induced artifacts and sample destabilization.

In addition, we elected to pair SurfACT with the SPT Labtech chameleon, a rapid blot-free plunge-freezing platform designed to minimize particle exposure to the AWI by dramatically reducing the time between sample deposition and vitrification. Beyond its ability to partially ‘outrun’ AWI interactions, the chameleon provides a high degree of automation and reproducibility in grid preparation, enabling systematic and user-independent control over ice thickness and quality.^7,32,33^ As demonstrated in our previous work, tuning ice thickness on the chameleon can be achieved in a predictable and programmable manner, decoupling specimen behavior from operator-dependent blotting variables and reducing a major source of variability in cryoEM sample preparation.^7^ We therefore leveraged the chameleon to isolate and evaluate the effects of SurfACT itself while minimizing confounding variability arising from ice thickness, blotting heterogeneity, or inconsistent vitrification conditions.^7,55^

### SurfACT Induces Comprehensive Hemagglutinin (HA) Viewing Distributions for Near-Atomic Resolution Reconstruction

Hemagglutinin (HA) is a homotrimeric glycoprotein of the influenza virus that has long been a biological target of interest for infectious diseases and is also the focus of numerous preferred orientation cryoEM studies.^35,56^ The HA trimer has an elongated geometry consisting of a globular head domain and stalk domain with a central three-fold symmetry axis. In the absence of any additives, HA micrographs are dominated by ‘top’ or ‘bottom’ views, looking down the C_3_ symmetry axis, with a noted absence of ‘tilt’ and ‘side’ views, resulting in poor map quality, stretched EM density, and limited stalk density (**Figure 2A-B, S2-6**).^35^ Diverse approaches have been used to recover views and isotropic density for the HA trimer, including tilted data collection, detergent addition, addition of conformationally-selective antibody fragments, data processing strategies, and/or machine-learning algorithms — all with limited success (**Supplementary Table 1**).^35,37–39^ As a result, HA remains a stringent benchmark for evaluating methods intended to overcome AWI-driven artifacts.

**Figure 2:**
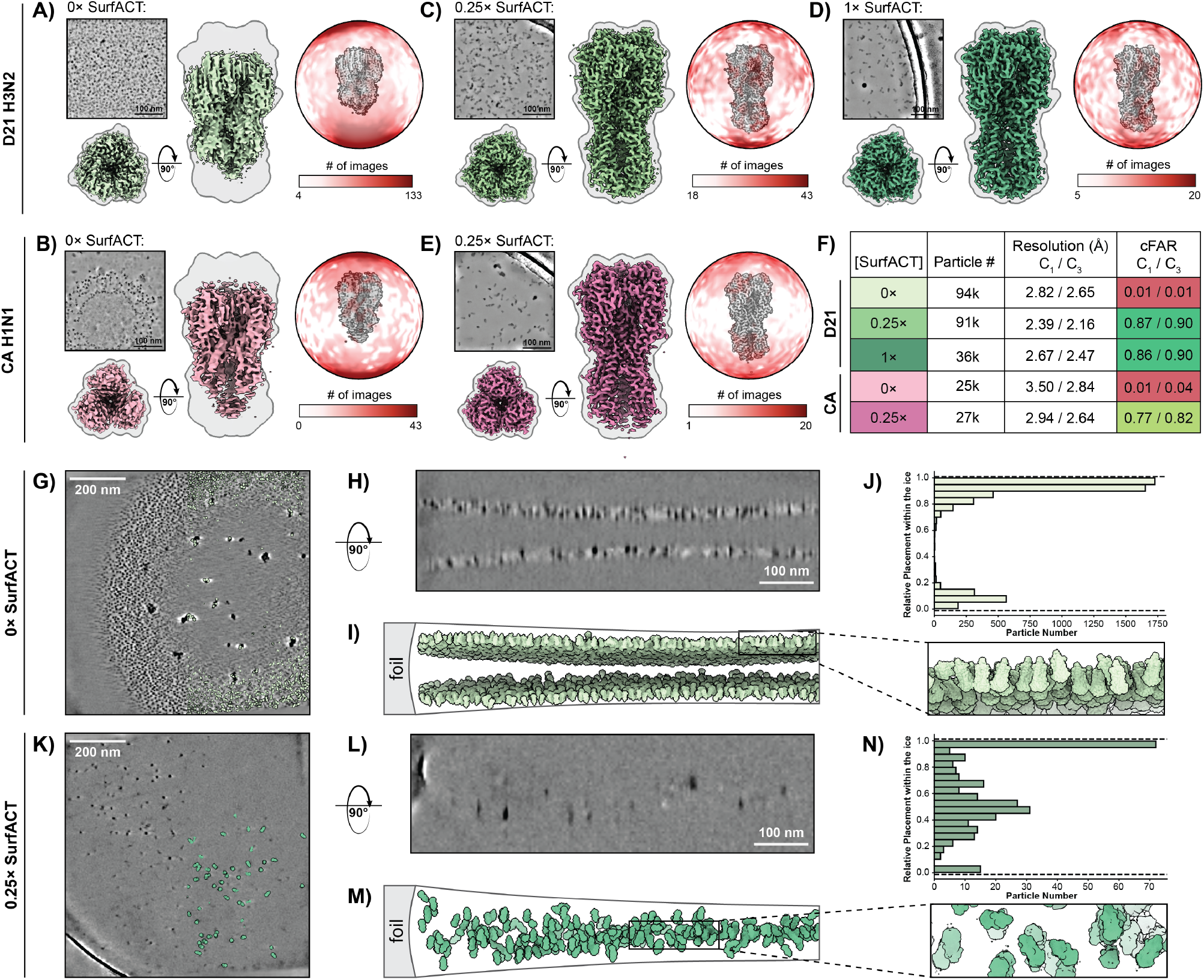
Hemagglutinin Preferred Orientation Mitigated Using SurfACT. **A-E**) HA cryoEM analysis: denoised micrograph, EM density in top-down and side views, and signal distribution shown as a heat map from white (low) to red (high). D21 H3N2 HA was assessed with 0× (**A**), 0.25× (**C**), and 1× (**D**) SurfACT. CA H1N1 HA was assessed with 0× (**B**) and 0.25× (**E**) SurfACT. An outline of intact HA density is shown. **F**) Table detailing particle number, resolution, and improved signal distribution (cFAR) value for final C_1_ and C_3_ symmetry reconstructions for all five HA datasets highlighting cFAR from SurfACT addition with similar or fewer contributing particles. cFAR values are colored from red (poor)–yellow–green (good). **G-J**) CryoET analysis of D21 H3N2 HA with 0× SurfACT. Representative tomogram slices in XY (**G**) and YZ (**H**) showing HA localized almost exclusively at the two AWIs. Tomogram reconstruction (**I**) with C3 cryoET HA volumes (light green; reconstructed from a 3-tomogram dataset) plotted on particle locations and showing orientations calculated from refined Euler angles. Particles adopt mostly uniform orientations pointing at the AWI, resulting in the ‘top’ or ‘bottom’ views seen by SPA (**I** *flyout*). **J**) Quantitative analysis of particle placement relative to tomogram ice boundaries by histogram. Particles were identified mostly near the ice boundaries with few to no particles identified in the center of the ice. **K-N**) CryoET analysis of D21 H3N2 HA with 0.25× SurfACT. Representative tomogram slices in XY (**K**) and YZ (**L**) showing HA distributed evenly. Tomogram reconstruction (**M**) with C_3_ cryoET HA volumes (darker green; reconstructed from a 12-tomogram dataset) show particles distributed throughout the ice cross-section in a variety of orientations (**M** *flyout*). **N**) Quantitative analysis by histogram shows particles were identified at the AWIs but majority of particles were distributed throughout the bulk ice.

To assess whether SurfACT could robustly address these challenges, we examined two antigenically and structurally distinct HA trimers — Darwin 2021 H3N2 (D21 H3N2) and California 2009 H1N1 (CA H1N1) — which both exhibit the canonical HA preferred orientation phenotype (**Figure 2A-B**).^57,58^ The impact from addition of SurfACT to both HA samples was immediately obvious, resulting in an abundance of newly accessed views, including tilt and side views, that can be easily observed by eye at the micrograph level, further exemplified in the 2-D class averages (**Figure 2C-E, S2-4**). Comparison of 3-D reconstructions from standard grid preparation (e.g., 0× SurfACT) and SurfACT-containing datasets demonstrates recovery of the stalk electron density, improved overall EM density without stretching artifacts, and higher nominal resolution (**Figure 2F**).

Although surfactants have been shown to improve particle behavior in cryoEM, their ability to alter surface tension modifies wicking behaviors and frequently leads to changes in particle density — namely the absence of particles at the center of the hole and clustering at the edge in areas of thicker ice, a phenomenon known as particle pushing.^59,60^ Brij and CHAPSO particularly contribute to particle pushing, and lower additive concentration overall is desired to prevent decreased particle concentrations often observed with these surfactants. To counter this phenomenon, we increased protein concentration, tested lower SurfACT concentrations, and collected closer to the edge of the grid hole (**Figure S7**). Our formulation working concentration (FWC) with all components 10-fold below CMC is defined as 1× SurfACT, and other concentrations tested are defined in relation to the working concentration (e.g. 0.25×). 0.25× SurfACT mitigated preferred orientation effects for HA and minimized changes to particle density in a sample that could not be highly concentrated, so this was used as the minimal SurfACT concentration tested throughout our study. With D21 H3N2 HA, SurfACT exhibits titratable effects for stronger and more complete stalk density as SurfACT concentration increases from 0.25× to 1×. The volume completeness and global resolution is starkly improved with 0.25× SurfACT, and EM density is further strengthened at the periphery of the stalk domain with the higher 1× SurfACT concentration (**Figure 2C-D**). In addition to qualitative assessment of the volume, we can evaluate the signal distribution in Fourier space that contributes to the reconstruction. Elevation plots provide a heatmap visual to illustrate signal distribution, where samples experiencing preferred orientation exhibit hotspots of signal in Fourier space corresponding to limited 2-D viewing angles and have regions with little or no signal (**Figure 2A-B**), while an ideal sample would have consistent relative signal across all regions of Fourier space (**Figure 2D**). Conical FSC area ratio (cFAR) is a metric of viewing distribution, with values from 0 to 1, where <0.5 indicates preferred orientation and closer to 1 indicates even signal distribution. Non-SurfACT HA datasets have extremely low cFAR (0.01) signifying pathological preferred orientation and large swaths of Fourier space devoid of any signal, despite moderately high FSC-estimated resolution values. With the lowest concentration of SurfACT tested, 0.25×, cFAR immediately jumps to ~0.8 for both CA H1N1 and D21 H3N2 HA trimers and further increases when C_3_ symmetry is imposed in refinement (**Figure 2F**). cFAR remains similarly high for D21 H3N2 HA with 1× SurfACT, but stronger particle pushing behavior caused the overall particle yield to be lower. With fewer than half as many particles as the 0.25× SurfACT dataset, the 1× SurfACT dataset has slightly lower nominal resolution (2.47 Å (C_3_) vs. 2.16 Å (C_3_)) but stronger and more continuous density at the edge of the stalk (**Figure 2D**).

### CryoET Demonstrates SurfACT Redistributes HA Particles from the AWI into Bulk Ice

To understand how the distribution of HA orientations is influenced by the AWI and changes upon introduction of SurfACT, we turned to cryogenic electron tomography (cryoET) to identify particle association with the AWI.^9,36,61,62^ Tomograms of the ice cross-section were reconstructed from tilt series data collections and, with the additional dimension (Z) afforded by cryoET, we can visualize where particles localize in the ice (**Figure S8**). In the absence of any surfactants, nearly all HA particles localized to the AWI (**Figure 2G-J**), while the addition of 0.25× SurfACT largely disrupted this interaction and redistributed particles to the bulk ice (**Figure 2K-N**). We performed template matching followed by subtomogram averaging (STA) to refine the positions and orientations of these particles, then mapped them back onto the tomograms using their refined Euler angles to investigate particle orientation — revealing that particles at the AWI are oriented perpendicular to the ice surface, providing the top-down view seen in SPA (**Figure 2I**), while particles in the bulk ice have random orientations and deliver varied 2-D views, including ‘tilted’ and ‘side’ views (**Figure 2M**). We also note that with the additional angles from tilt series data collection and STA, cryoET HA reconstructions with 0× Sur-fACT feature a complete stalk domain (**Figure 2I**) compared to SPA reconstructions (**Figure 2A**), indicating the AWI particle association only biases particle viewing distribution and does not induce other artifacts like denaturation.

### SurfACT Rescues the αIII Domain of MoFeP to Improve Map Isotropy and Quality

Molybdenum-iron protein (MoFeP) from the *Azotobacter vinelandii* nitrogenase enzyme has been shown to similarly suffer from interaction with the AWI when not in complex with other nitrogenase subunits.^63^ While views are not as dramatically limited as the single top-down HA view, minimal MoFeP viewing distribution results in loss of electron density for the dynamic *α*III domain, which helps encase the catalytically important iron-molybdenum cofactor (FeMoco) (**Figure 3A, S9-10**). Visualization of *α*III and FeMoco is critical for the study of catalytic states for nitrogenase, whose nitrogen fixation capabilities sustain productivity in all ecosystems, and has been shown to be dynamically coupled to catalytic state.^63–65^

**Figure 3:**
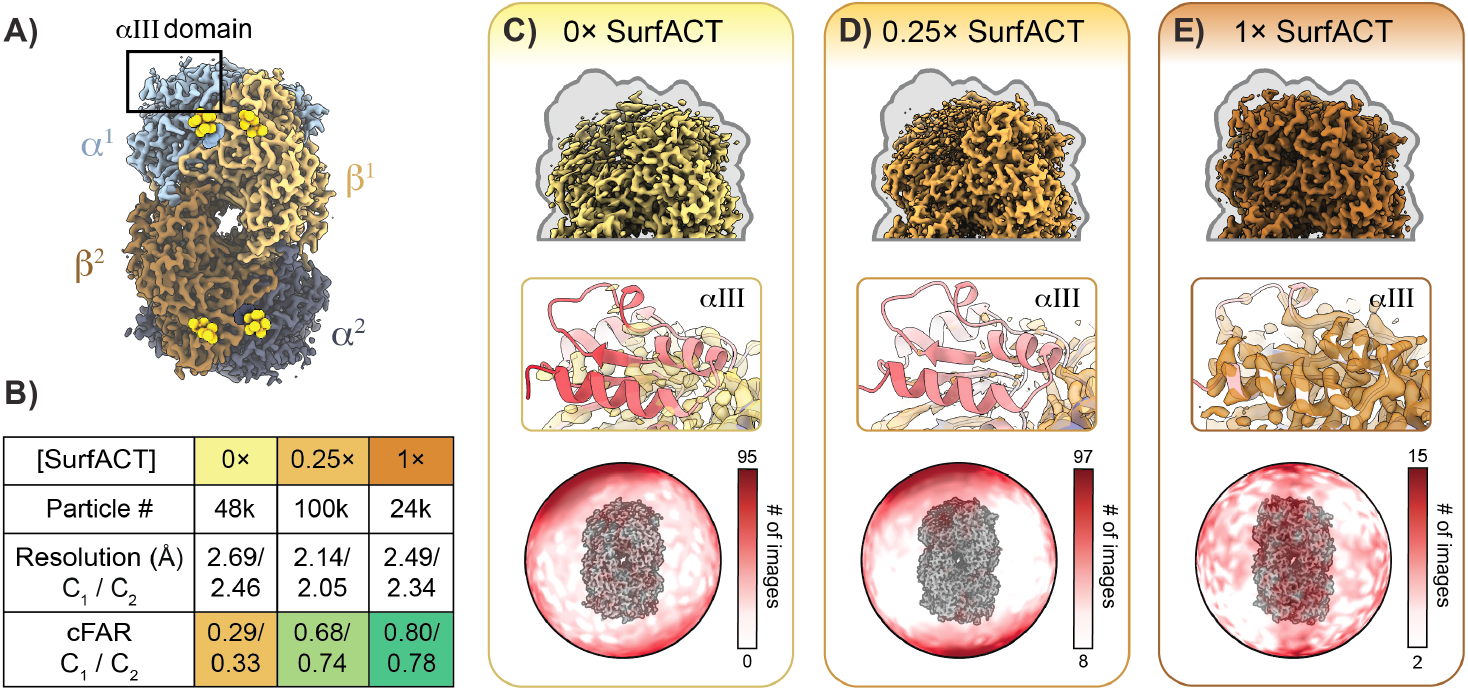
SurfACT Recovers MoFeP’s *α*III Domain and Improves Map Quality. **A**) MoFeP colored by chain with metal clusters shown in yellow. The *α*III domain, which can be lost due to AWI-induced artifacts, is highlighted. **B**) Statistics for final EM densities across SurfACT concentrations highlighting that SurfACT datasets have improved resolution and signal distribution even with significantly lower particle count than 0× SurfACT. cFAR values are colored from red (poor)-yellow-green (good). **C-E**) EM density of MoFeP, density-model fit for peripheral helix in *α*III domain with corresponding models colored by B-factor blue (low)-white-red (high), and signal distribution shown as a heat map from white (low) to red (high). An outline of intact MoFeP density is shown.

Again, EM grid preparation with SurfACT addition improves resolution and signal distribution of resulting EM densities without reliance on enforced symmetry (**Figure 3B**). Without SurfACT, there is near complete loss of the dynamic *α*III domain, stretching artifacts of adjacent EM density, and a poor cFAR of 0.29 (**Figure 3B-C**). Addition of 0.25× SurfACT yields some improvements for MoFeP, but not quite as potent of an effect as it had for the HA trimer.

Some density is recovered in *α*III and cFAR rises to 0.68, but the volume remains incomplete, even at a nominal 2.14 Å (C_1_ symmetry) resolution (**Figure 3B, 3D**). With 1× SurfACT, the *α*III domain is recovered completely (**Figure 3E**). cFAR reaches 0.80, indicating well-distributed signal in Fourier space, eliminating AWI-induced issues in the final volume, and improving volume quality and completeness while increasing nominal resolution from 2.69 Å (C_1_ symmetry) to 2.49 Å (C_1_ symmetry) with less than half the particle count (**Figure 3B, 3E**).

### SurfACT Suppresses the Aldolase “Broken Protomer” Phenotype

Rabbit muscle aldolase has also been involved in many cryoEM studies to evaluate microscope and hardware capabilities, data collection and processing strategies, and cryoEM sample preparation.^66–71^ Despite aldolase’s utility as a cryoEM test specimen, the homotetramer exhibits persistent loss of EM density at the periphery of one protomer and, as a result, symmetry has been frequently enforced in studies evaluating cryoEM sample preparation and data processing strategies. In datasets prepared without surfactant additives (0× SurfACT), we observed this same “broken protomer” phenotype in our ~2.42 Å reconstruction (C_1_ symmetry), with approximately half of one subunit failing to resolve continuous density despite high nominal resolution (**Figure 4A, S11-14**). This phenotype has previously been addressed by larger data collections at high particle concentration, meticulous particle pruning, and reliance on enforced symmetry in data processing.^66,70^ Here, we address the root cause of the missing density - missing signal in the micrographs from sample destabilization, instead of artificially filling in information with enforced symmetry.

**Figure 4:**
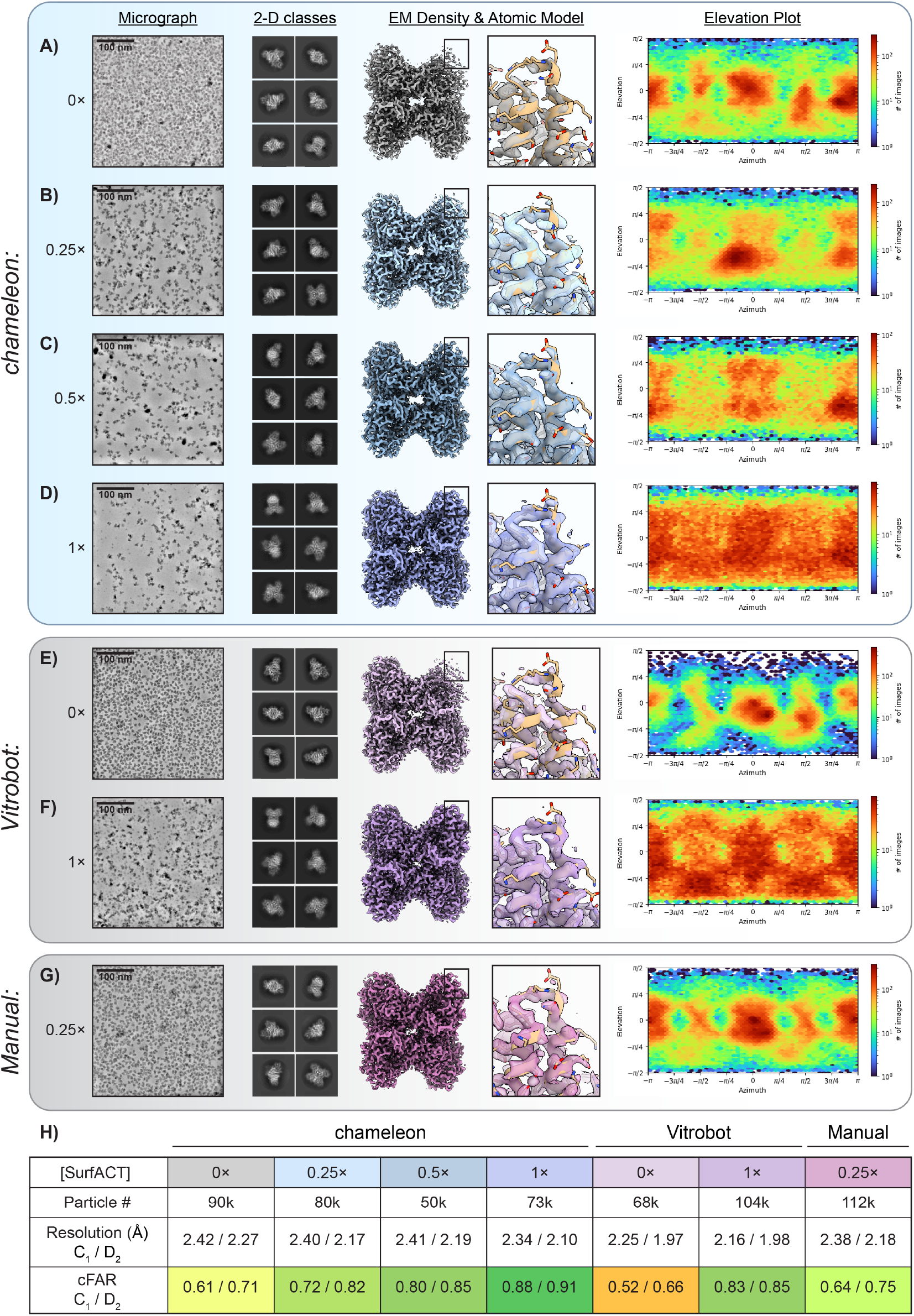
SurfACT Prevents Loss of Aldolase Protomer Density and Improves Viewing Distribution Across Diverse Freezing Modalities. Thorough survey of SurfACT concentrations and freezing modalities: chameleon (**A-D**), Vitrobot (**E-F**), and manually operated plunge freezing (**G**). For each dataset, a representative denoised micrograph, representative 2-D classes ordered by particle number, refined EM density (C_1_ symmetry), flyout of model-to-density agreement of a peripheral *α*-helix of the destabilized protomer, and cryoSPARC elevation plot to demonstrate signal distribution are provided. **H**) Statistics for final EM densities across SurfACT concentrations highlighting that SurfACT datasets have improved resolution and signal distribution. cFAR values are colored from red (poor)-yellow-green (good).

To test this directly, we conducted a systematic review of SurfACT concentrations, across a range of 0×, 0.25×, 0.5×, and 1×, and grid freezing techniques with aldolase. Data quality for aldolase gradually improves with addition of SurfACT, including observation of additional 2-D views, notably a “butterfly” view with all four intact protomers clearly visible (**Figure 4B**), and more particles have three or four protomers visible in 2-D classes from higher (0.5×, 1×) Sur-fACT datasets (**Figure 4C, 4D**). There is a gradient of aldolase volume completeness and signal distribution evidenced by elevation plots and cFAR values across the datasets from 0× to 1× SurfACT. In the dataset containing 1× SurfACT, the broken protomer is fully recovered with continuous EM density in the distal *α*-helices, and cFAR is correspondingly improved from 0.42 (without SurfACT) to 0.87 (with 1× SurfACT) (**Figure 4D**). It’s important to note that the nominal resolution for 0× and 1× Sur-fACT aldolase is nearly identical (2.41 and 2.40 Å; C_1_ symmetry) while the volume quality is drastically improved in the 1× SurfACT dataset (**Figure 4A, D**). Nominal resolution is a poor indicator of data quality, so we largely focus on other measures like map completeness and viewing distribution.

### SurfACT Improves Sample Behavior Using Both Traditional Blot-And-Plunge and Blot-Free Vitrification Protocols

To expand utility of SurfACT beyond the blot-free chameleon, we tested traditional blot-and-plunge freezing methodologies using both the Vitrobot Mark IV grid preparation device and a manually operated grid-freezing device.^6,7^ Aldolase on Vitrobot-prepared grids, without or with 1× SurfACT, behaved very similarly to chameleon datasets, with comparable observed viewing distribution and final volume quality (**Figure 4E-4F, S15-16**). We were able to obtain a 2.16 Å resolution (C_1_ symmetry) 3-D reconstruction of intact aldolase using 1× SurfACT (**Figure 4F**), further improved to 1.98 Å resolution with D_2_ symmetry enforced (**Figure 4H**).

We postulated that the higher resolution Vitrobot-frozen 1× SurfACT dataset (2.16 Å; C_1_ symmetry) compared to the chameleon-frozen 1× SurfACT dataset (2.34 Å; C_1_ symmetry) is likely a result of more rigid sample supports, namely, the gold UltrAuFoil grids available for Vitrobot use versus traditional carbon foil grids (with nanowires) used for chameleon grid freezing. Since gold grids have been demonstrated to yield less beam-induced motion compared to copper-carbon grid combinations, which could result in higher resolution 3-D reconstructions,^72–74^ we wanted to test if the resolution difference between the Vitrobot and chameleon datasets was due solely to the grid composition or a difference in SurfACT behavior across freezing methods.

We froze aldolase sample with 1× SurfACT on gold-coated chameleon grids and were able to obtain a 2.43 Å resolution reconstruction (C_1_ symmetry) with <50% of the particle count of the Vitrobot 1× dataset (34k vs. 83k, respectively). D_2_-symmetrized reconstruction went to 2.17 Å, compared to 2.16 Å D_2_ reconstruction obtained using the Vitrobot, and cFAR values were nearly identical (0.86 vs 0.85) for both datasets (**Figure S17-19, 4F, 4H**). Consequently, we conclude that differences in grid composition and particle count are responsible for discernable differences between SurfACT datasets across freezing methods and do not limit application of SurfACT to one modality over another. Additionally, compared to chameleon-prepared samples, SurfACT performed equally well with aldolase on grids frozen using a manually operated plunging device (**Figure 4G-H, S20-21**). Manual-plunge frozen aldolase with 0.25× SurfACT achieved volume completeness similar to chameleon-frozen 0.5× and signal distribution comparable to chameleon 0.25×. Through this systematic aldolase screen, we demonstrate that SurfACT performs consistently to ameliorate AWI-induced artifacts across all commonly utilized cryoEM grid freezing modalities.

### Accommodating SurfACT Changes to Particle Density and Distribution During Imaging

Through this thorough screen of aldolase against Sur-fACT concentrations and freezing modalities, we additionally evaluated documented surfactant-induced “particle pushing” behavior attributed to SurfACT. All SurfACT samples were prepared with at least twice the protein concentration compared to non-SurfACT samples. For chameleon samples, the 0× SurfACT was made with 6 mg/ml aldolase, and all SurfACT-containing samples were prepared at 12 mg/ml aldolase. The Vitrobot samples were prepared at 1 mg/ml aldolase for 0× SurfACT and 4 mg/ml for 1× SurfACT. As a baseline, we observe much lower particle concentration with any amount of SurfACT added, requiring the higher initial protein concentration (**Figure 4**). At 0.25× SurfACT, particles are present across all regions of the grid hole, including the center, and particle density is relatively uniform across the hole (**Figure S7**). At 0.5× SurfACT, a gradient of particle concentration emerges, and particles are sparser in the center of the hole (**Figure S7**). At 1× SurfACT, the gradient intensifies with very few particles in the center, requiring imaging closer to the edge of the hole (**Figure S7**). We advise imaging in the middle region of the concentration gradient when using SurfACT, in between the center and edge of the hole, to balance particle concentration and ice thickness for optimal data collections. Despite lower particle concentration with addition of SurfACT, particles are intact and provide more varied views, increasing yield in particle pruning and requiring fewer particles for a complete 3-D reconstruction.

### CryoET Confirms Aldolase Protomer Loss is Due to Denaturation at AWI

We performed a similar template matching and STA pipeline to aldolase particles visible in our tomograms, and as with the HA trimer, aldolase particles are found almost exclusively at the AWI in the absence of SurfACT (**Figure 5A, S8, S22**). With addition of 0.25× SurfACT, a similar proportion of particles associate with the AWI (~90%)(**Figure 5B, S22**) but, 1× SurfACT nearly eliminates AWI interactions and over 80% of particles localize to the bulk ice (**Figure 5C, S22**). Even with the side views afforded by a −54° and +54° tilt series, in datasets with a large portion of AWI particles (i.e. 0× and 0.25×), one protomer remains broken, and the viewing distribution orients through the broken protomer (**Figure 5D**). To probe this missing density, we separated the 0.25× SurfACT particle stack into groups based on each particle’s proximity to the AWI (**Figure 5E**). Particles within 100 Å of the upper and lower ice boundary were grouped as ‘top AWI’ and ‘bottom AWI’, respectively, and reconstructed separately, and EM density loss persists with dominant preferred orientation viewed through the broken protomer (**Figure 5F-G**). Consistent with this observation, when we back-project the particle orientations onto the tomogram using their refined Euler angles calculated in the refinement step, the broken protomers all point toward the AWI (**Figure 5F-G**). Due to the nature of tomographic reconstructions, a more complete volume should be obtained if the interaction of aldolase with the AWI was solely due to missing views (see **Figure 2I**). However, the information is missing entirely, which allows us to conclude that the broken protomer is a product of protein denaturation at the AWI. When particles from the center or bulk ice are segregated from AWI particles and reconstructed alone, the resulting volume is a complete tetramer with well-dispersed signal (**Figure 5H**). Thus, SurfACT addition circumvents the AWI interaction that partially denatures aldolase and recovers views that are inaccessible by traditional single-particle data collection.

**Figure 5:**
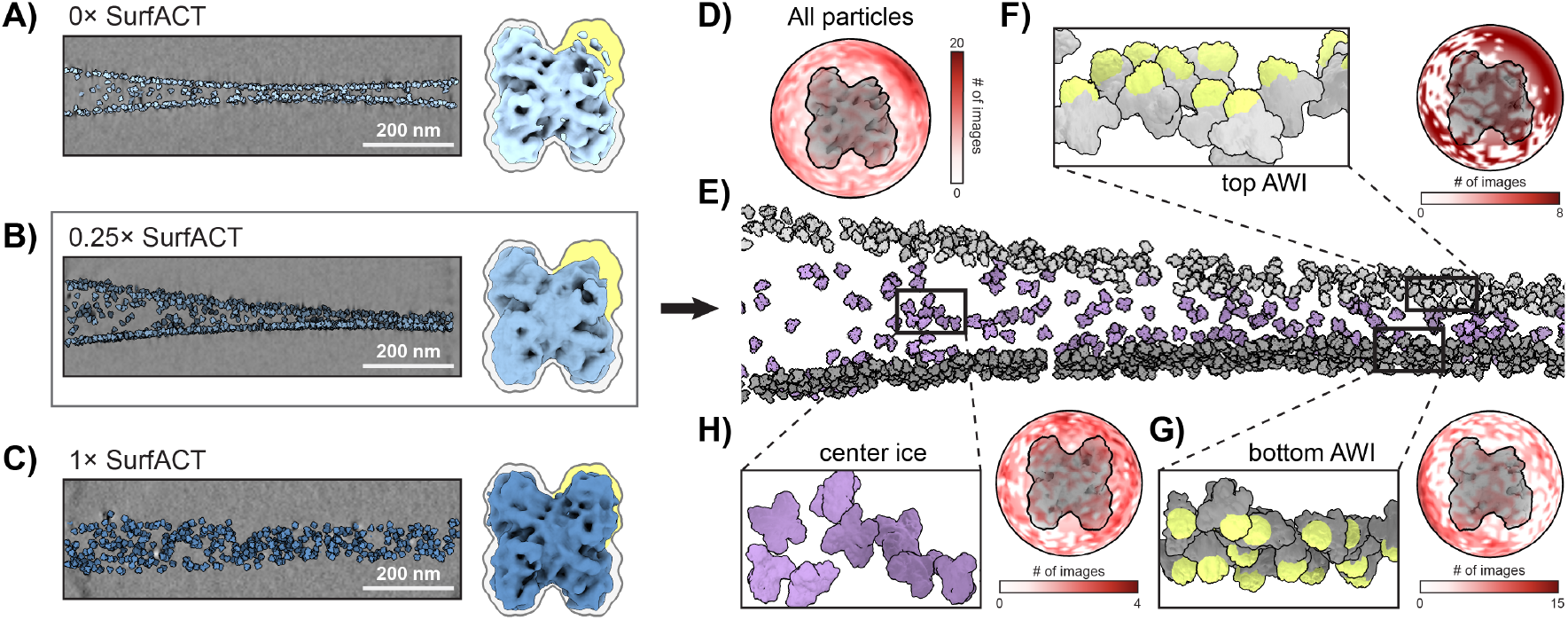
SurfACT Prevents Loss of Aldolase Protomer Density and Improves Viewing Distribution Across Diverse Freezing Modalities. CryoET analysis of aldolase across SurfACT concentrations. **A-C**) YZ representative tomogram slice shows particles are redistributed from the AWI to the bulk ice with increasing SurfACT concentration. Correlating STA densities from **A, B**, and **C**, respectively, are shown on the right with an outline of intact aldolase shown. Area highlighted in yellow indicates broken protomer phenotype. **D**) Signal distribution of the cryoET reconstruction obtained using all particles from the 0.25× SurfACT dataset. E) Aldolase STA reconstructions with 0.25× SurfACT grouped by location at the top AWI (*light grey*), bottom AWI (*dark grey*), and center ice (*purple*). Top and bottom AWI groups included all particles within 100 Å of the ice boundary. **F-G**) Top (**F**) and bottom (**G**) AWI particles have similar orientations with one protomer pointing at the AWI. Reconstructions of each particle stack results in missing EM density and a hotspot in signal distribution. **H**) Center ice particles provide more views for a complete aldolase reconstruction and more disperse signal distribution.

### “Particle Pushing” is the Absence of an On-Grid Concentrative Effect

Lastly, we used cryoET to investigate the particle pushing behavior attributed to SurfACT and surfactant use generally. By calculating the volume of the ice cross-section and relating the number of particle picks per tomogram, we can calculate the final on-grid protein concentration. Without SurfACT, aldolase is ~ 2.3-fold the concentration applied to the grid, and particles at the AWIs, within 100 Å of the ice boundaries, were found to be ~2.9-fold the applied aldolase concentration (**Figure S23**). This provides direct evidence for a concentrative effect occurring at the AWI during the blotting/wicking process. 0.25× SurfACT tomograms contain ~0.9-fold aldolase particles relative to applied concentration, with ~1.2-fold aldolase concentrated at the AWIs (**Figure S23**). Particles remain localized at the AWI with 0.25× SurfACT, but the concentrative effect is diminished in overall tomogram concentration, from ~2.3-fold to ~0.9-fold applied sample, and concentration at the AWIs, from ~2.9-fold to ~1.2-fold. In 1× SurfACT tomograms, the AWIs were not marked with aldolase particles, so we could not reliably define the ice boundary for mask generation. The center ice particle concentration, where we are confident there are no empty voxels, is ~0.6-fold the applied aldolase concentration (**Figure S23**). While particle pushing is present, evidenced by the concentration gradient across the hole, the phenomenon is minimally responsible for the higher sample requirements with SurfACT. Surfactant presence slightly decreases the effective concentration of protein in the grid hole (~0.6-0.9-fold), but the particle “dilution” observation is primarily driven by the absence of AWI interactions concentrating the protein (~2.3-fold) during wicking, making the comparison with and without additives more drastic.

## Discussion

Our systematic survey of surfactant use across deposited cryoEM structures reveals a field dominated by empirical, single-additive strategies that yield sporadic, sample-specific successes, which do not generalize across samples or preparation workflows. Despite widespread recognition that AWI interactions underlie many cases of preferred orientation and structural degradation, no single additive has emerged as a universally effective solution, and usage in the field remains variable and often trend-driven (**Figure 1, S24**). We developed SurfACT as a rationally designed, low concentration surfactant cocktail intended to balance complementary interfacial behaviors while minimizing the destabilizing and stochastic effects often encountered with individual surfactants used in isolation.

Across three structurally and functionally distinct protein systems — hemagglutinin (HA), molybdenum-iron protein (MoFeP), and aldolase — our results demonstrate that Sur-fACT reproducibly mitigates AWI-driven artifacts, restores missing structural information, and improves viewing distribution by redistributing particles from the AWI into the bulk vitreous ice. SurfACT consistently and significantly improves data quality for four distinct samples, with reconstructions approaching 2 Å resolution or better and cFAR values 0.8 or higher without necessitating symmetry for signal recovery.

For the HA trimer examples, 2-D class averages were almost entirely dominated by a single “top-down” view without SurfACT, but with SurfACT we access side and tilt views necessary for complete HA reconstruction and visualization of the therapeutically important stalk domain (**Figure 2**). SurfACT was highly effective at low concentration (0.25×) with HA from two different flu strains, a promising indicator of compatibility across other variants. Similar improvements were also observed with MoFeP. Namely, without additives, the dynamic *α*III domain of MoFeP could not reproducibly be visualized by cryoEM without significant sorting or imposed symmetry. With 1× SurfACT, MoFeP viewing distribution was recovered to reconstruct the complete *α*III domain and visualize the N_2_ reduction active site, allowing for future study of MoFeP catalytic states (**Figure 3**). Finally, SurfACT recovers a destabilized aldolase subunit by disrupting AWI interactions that partially denature the protein and additionally improves viewing distributions (**Figure 4**). We largely focus on measures like viewing distribution, map completeness, and local resolution in our assessment of SurfACT effects since global resolution is a poor indicator of data quality and can conceal notable domain weaknesses.

SurfACT aims to disrupt the root cause of preferred orientation in cryoEM — AWI interactions during grid preparation. By reducing protein-AWI interactions, more molecules tumble randomly in the bulk ice showing unique 2-D views, meaning the same number of particles hold more information in data analysis (**Figure 6**). We demonstrate in the STA reconstructions of D21 H3N2 HA and aldolase that particles from the bulk ice provide isotropic signal distribution and that maximizing the proportion of bulk ice versus AWI-localized particles with SurfACT is an effective strategy to improve cryoEM volumes (**Figures 2 and 5**).

**Figure 6:**
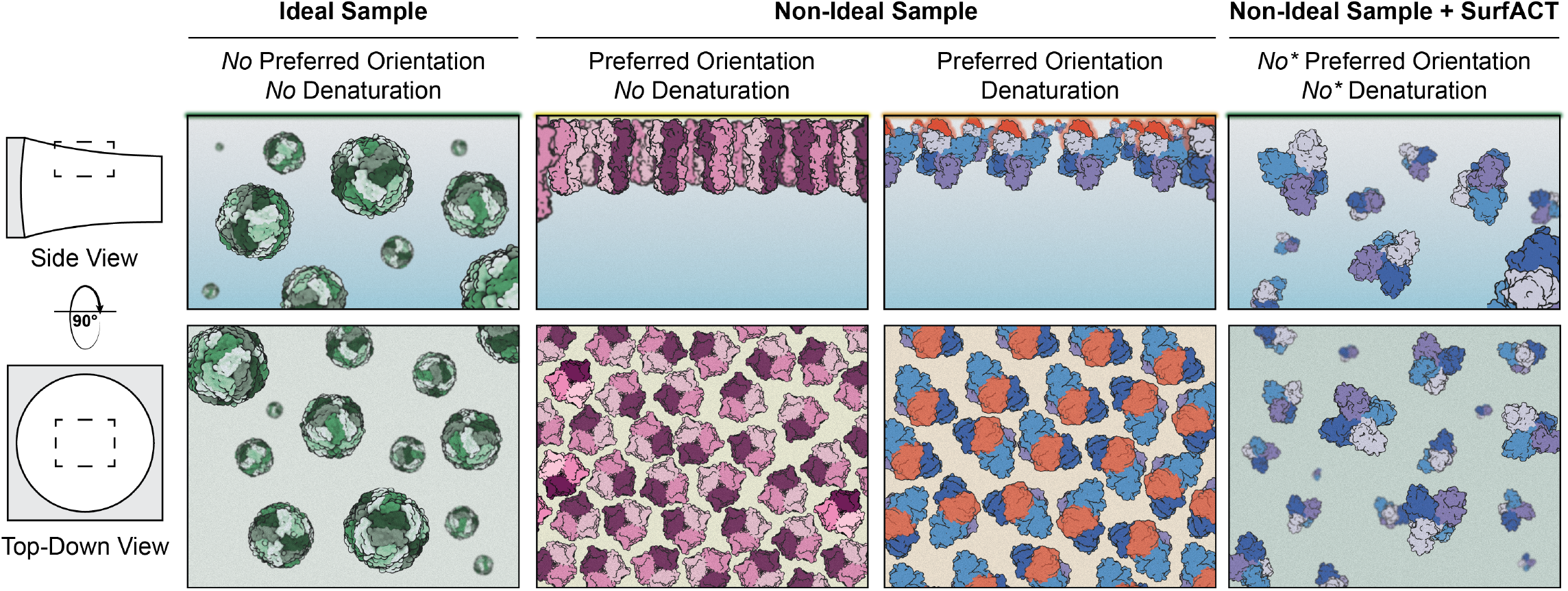
AWI Interactions Ameliorated by SurfACT. With ideally behaving EM samples, like apoferritin (*green*), particles tumble in bulk solution and randomly orient in vitreous ice. Under non-ideal conditions, many samples localize to the AWI during grid freezing and experience AWI-induced artifacts. HA (*pink*) experiences pathological preferred orientation but no AWI-induced denaturation, so missing SPA density can be recovered by STA. Aldolase (*blue* with broken protomer colored *red*) has preferred orientation but is also denatured at the AWI, so tilted-data collection cannot fill in missing EM density. Addition of SurfACT eliminates AWI interactions resulting in improved distribution and sample recovery. ^∗^With 1× SurfACT, aldolase (*blue*) SPA reconstructions produce near-perfect cFAR (0.88) and high-quality volumes with no stretching or other artifacts of preferred orientation (see **Figure 4**), and cryoET reconstruction produces a complete volume, indicating intact, non-denatured aldolase (see **Figure 5**).

SurfACT was demonstrated to be effective across numerous freezing methods, including the blot-free SPT Labtech chameleon, traditional blot-and-plunge Vitrobot, and manually operated blot-and-plunge freezing, and provides high-quality data that minimizes or eliminates reliance on enforced symmetry in data processing to obtain an isotropic, complete 3-D reconstruction. Although our formulation and FWC of 1× SurfACT was shown to be effective for all samples evaluated, we note that if a sample cannot be highly concentrated, 0.25× SurfACT provides notable volume improvements with mild particle pushing and higher particle concentration compared to 1×. Some proteins, like HA, respond strongly to 0.25× SurfACT, so this may be sufficient. If more surfactant is required, like MoFeP and aldolase, 1× SurfACT further improves viewing distribution and volume completeness, but larger datasets may be necessary to account for perceived lower particle density since the concentrative AWI effect does not inflate apparent sample concentration. Due to the physicochemical effects of surfactants, changes in surface tension may require selective imaging in slightly thicker ice closer to the edge of the grid hole, and considerations around SurfACT concentrations, protein concentration, and ice thickness must be simultaneously valued. We additionally tested SurfACT on human hemoglobin, which does not experience preferred orientation, and validated that there were no deleterious effects beyond micrograph density and particle pushing (data not shown).

With all four samples tested, we prioritized generation of volumes without enforced symmetry to accurately evaluate AWI-induced damage. In the case of aldolase, we utilized symmetrized volumes as a 3-D sorting tool in data processing. Specifically, a non-uniform 3-D refinement was performed with either C_1_ or D_2_ symmetry, and the resulting volumes were used for heterogeneous refinement to sort ‘intact’ versus ‘broken’ aldolase particles (**Figure S11**,**13**,**15**,**18**,**20**). We employed this particle-pruning strategy across all aldolase datasets and observed a correlation between the proportion of ‘intact’ particles and SurfACT concentration. In general, we observed increasing percentages of ‘intact’ aldolase particles as a function of [SurfACT], from 36% in the absence of surfactant to *>*64% with 1× SurfACT, indicating that a larger portion of particles have information for all four aldolase protomers with increased SurfACT concentration. This data correlates with the proportion of AWI-associated particles we see by cryoET, which do not experience AWI-induced denaturation and provide isotropic signal distribution (**Figure 5**).

Through a combination of SPA and cryoET, we provide a comprehensive analysis of the SurfACT sample additive and its suitability and versatility across different protein classes, sizes, and complexities. SurfACT demonstrates a dramatic recovery in signal distribution and map completeness that we can directly attribute to the proportion of bulk ice particles, as measured by cryoET. We also provide evidence of the novel finding that particles are concentrated at the AWI during grid preparation, by calculating concentration of particles relative to the volume of the tomogram ice cross-section. SurfACT tackles the pervasive problem of preferred orientation by improving efficacy in grid preparation - refining samples previously considered difficult or unsuitable for cryoEM.

## Methods

### Surfactant Survey Across CryoEM Publications

To generate an initial list of research papers that utilize surfactants to overcome preferred orientation, two different programmatic search and screen methods were employed. The first utilized EBI web services (“https://www.ebi.ac.uk/europepmc/webservices/rest/search”) where full-text articles were screened for keywords indicative of surfactant inclusion during cryoEM sample preparation. Keywords include a surfactant (“OG”, “Octyl *β*-D-glucoside”, “Octyl beta-D-glucoside”, “OBG”, “Octyl *β*-glucoside”, “Octyl beta-glucoside”, “beta-octyl glucoside”, “LMNG”, “Lauryl Maltose Neopentyl Glycol”, “CHAPS”, “CHAPSO”, “Brij-35”, “brij-35”, “FOM”, “fluorinated octyl maltoside”, “A8-35”, “amphipol A8-35”, “A835”, “NP40”, “NP-40”) and cryoEM (“cryoEM”, “cryo-EM”, “cryo electron microscopy”) and an associated EMDB entry (“EMD”, “EMD-”). An example can be found in **Supplementary Table 1**. This generated a list of 2,479 entries that contained keywords for surfactant addition during cryoEM sample preparation. The second method employed the EMDB, where a list of all EMDB codes were generated using (“https://www.ebi.ac.uk/ebisearch/ws/rest/emdb”). Then, the EMDB codes were filtered to generate a list of unique DOIs, which was 11,328 entries. Using (“https://api.unpaywall.org/v2/“) full-text publications were screened using the keywords for both. This generated a list of 1,604 articles. Entries we did not have access to were not included. The two lists were combined, and duplicates were removed to generate a final list of 2,818 entries.

All 2,818 research papers listed were manually screened, and surfactants added immediately prior to grid preparation were deemed “hits”. If the surfactant was used for protein solubilization or an earlier protein purification step not consistent with EM grid preparation, the entry was discarded. This resulted in a final list of 622 publications that detailed the use of surfactants for preferred orientation and/or sample destabilization. To ensure accuracy, we used a “few shot” approach using a LLM (Large Language Model) to validate our list, where both positive and negative examples were provided.

### Preparation of the SurfACT Mixture

A 20× SurfACT solution was prepared of 1% (w/v) 3-([3-cholamidopropyl]-dimethylammonio)-2-hydroxy-1-propanesulfonate (CHAPSO; Anatrace), 0.096% (w/v) 1H, 1H, 2H, 2H-perfluorooctyl)-*β*-D-maltopyranoside (FOM; Anatrace), 5% (w/v) amphipol A8-35 (Anatrace), and 1.2% (w/v) Brij-35 (Anatrace), stored in aliquots at −20°C. 20× SurfACT was diluted to 2×, 1×, or 0.5× in the freezing buffer for each sample and mixed 1:1 (v/v) gently with protein sample immediately prior to EM grid preparation. The final concentrations for 1× SurfACT (FWC) are 0.05% (w/v) CHAPSO, 0.0048% (w/v) FOM, 0.25% (w/v) amphipol A8-35, and 0.06% (w/v) Brij-35. All components are 10-fold below their critical micelle concentration (CMC) or, in the case of A8-35 which does not form traditional micelles, approximate protein solubilizing concentration (**Figure S1**).

### Preparation of Hemagglutinin (HA) Trimer Samples for CryoEM Analysis

Strain A/California/4/2009 H1N1 with E47K HA2 stabilizing mutation (CA H1N1) and Strain A/Darwin/6/2021 H3N2 (D21 H3N2) HA samples were kindly provided at 6.2 *µ*M and 39 *µ*M, respectively, by the Ward group at Scripps Research and used without modification unless specified. 0× SurfACT CA H1N1 HA was prepared at 6.2 *µ*M. 0.25× SurfACT CA H1N1 HA sample was concentrated to 11.3 *µ*M and mixed 7:1 (v/v) with 2× SurfACT for a final concentration of 9.9 *µ*M. 0× SurfACT D21 H3N2 HA was prepared at 10.5 *µ*M. 0.25× and 1× SurfACT samples were mixed 1:1 (v/v) with SurfACT for a final concentration of 19.5 *µ*M.

### Preparation of Azotobacter vinelandii Molybdenum Iron Protein (MoFeP) for CryoEM Analysis

Preparation of wild-type *Azotobacter vinelandii* molybdenum iron protein (MoFeP) (*Av* strain DJ200) was performed as previously detailed, and flash frozen under liquid nitrogen until use.^75^ MoFeP without SurfACT was prepared at 5 *µ*M. 0.25× and 1× SurfACT MoFeP samples were prepared at 30 *µ*M and mixed 1:1 (v/v) with 0.5× and 2× SurfACT, respectively, for a final MoFeP concentration of 15 *µ*M.

### Preparation of Rabbit Muscle Aldolase for CryoEM Analysis

As previously described, lyophilized rabbit muscle aldolase (Sigma-Aldrich A2714) was resuspended in 10 mM HEPES (pH 8) and 50 mM NaCl and centrifuged at 20,000× g for 10 min at 4°C prior to diluting to the desired concentration.^66^ Aldolase samples without SurfACT were prepared at 6 mg/ml for chameleon freezing and 1 mg/ml for Vitrobot freezing. All SurfACT samples were prepared at 2× protein concentration to be mixed 1:1 (v/v) with 0.5×, 1×, or 2× SurfACT for a final concentration of 0.25×, 0.5×, or 1× SurfACT. SurfACT samples had a final aldolase concentration of 12 mg/ml for chameleon freezing, 4 mg/ml for Vitrobot freezing, and 6 mg/ml for manually operated plunge freezing.

### CryoEM Grid Preparation

The SPT Labtech chameleon was prepared according to established protocols.^7^ Liquid ethane was maintained within a temperature range of −173 to −175°C, and the humidity shroud was maintained at *>*75% relative humidity during grid vitrification. For each sample, 2-3 chameleon self-wicking grids (Quantifoil Active 300 mesh; Quantifoil) were loaded into the instrument and glow discharged for 20 seconds at 12 mA. 10 *µ*L of sample was loaded into the sample vial and 7 *µ*L was aspirated into the dispenser and subsequently tested for proper dispensing within the chameleon software before sample deposition onto the grid. MoFeP sample vials were prepared for minimal oxygen exposure as described previously.^55^ Once ideal sample dispensing conditions were obtained, grids were frozen with 600-650 ms plunge time. Grids were stored under liquid nitrogen until needed.

Preparation of aldolase grids using a Vitrobot Mark IV (Thermo Fisher Scientific) were performed similarly using described protocols.^76^ Briefly, 3.2-3.4 *µ*L of aldolase sample with 0× or 1× SurfACT were applied directly to an UltraAuFoil R1.2/1.3 (Quantifoil) grids that had been pre-treated using a Solarus II (Gatan) plasma cleaner. Grids were stored under liquid nitrogen until needed.

Manual-plunge frozen aldolase grids were prepared on UltraAuFoil R1.2/1.3 (Quantifoil) grids that had been freshly plasma-cleaned using a Solarus II (Gatan) plasma cleaner. Grids were prepared using a custom manually operated plunge freezer designed by the Herzik Lab located in a humidified (*>*95% relative humidity) cold room (4°C).^6,70^ 3 *µ*L of the sample was applied to the grid surface followed by manual blotting for ~5-6 s using Whatman No. 1 filter paper before vitrifying in a 50% ethane/50% propane liquid mixture cooled by liquid N2.^6,70^ Grids were stored under liquid nitrogen until data collection.

### Single-Particle CryoEM Data Collection

All data was acquired at UCSD s CryoEM Facility using a Titan Krios G4 (Thermo Fisher Scientific) operating at 300 kV equipped with a Selectris-X energy filter (Thermo Fisher Scientific). All images were collected at a nominal magnification of 165,000× in EF-TEM mode (with a calibrated pixel size of 0.735 Å) on a Falcon4 direct electron detector (Thermo Fisher Scientific) using a 10 eV slit width. Micrographs were acquired in electron-event representation (EER) format with a cumulative electron exposure of 60 electrons/Å^2^ with a nominal defocus range of *−*1 to *−*2.0 *µ*m. Data were collected automatically using EPU2 (Thermo Fisher Scientific) with aberration-free image shift. Data was monitored during collection using cryoSPARC Live (Structura Bio) where movies were patch motion corrected and patch CTF estimated on-the-fly. HA datasets: D21 H3N2 HA 0×, 0.25×, and 1× SurfACT datasets contained 500, 1500, and 1482 micrographs, respectively (**Supplementary Table 2**). CA H1N1 HA 0× and 0.25× SurfACT datasets contained 458 and 2000 micrographs, respectively (**Supplementary Table 3**). MoFeP datasets: 0×, 0.25×, and 1× SurfACT MoFeP datasets contained 1010, 1401, and 1309 micrographs, respectively (**Supplementary Table 5**). Aldolase datasets: 0×, 0.25×, 0.5×, and 1× Sur-fACT aldolase chameleon-frozen datasets contained 1137, 1057, 1055, and 1029 micrographs, respectively. The 1× SurfACT aldolase dataset frozen on chameleon gold-coated grids contained 863 micrographs. Vitrobot-frozen 0× and 1× SurfACT aldolase datasets contained 476 and 1707 micrographs, respectively. The manual-plunge frozen 0.25× SurfACT aldolase dataset contained 1345 micrographs (**Supplementary Table 6-8**).

### Single-Particle CryoEM Data Processing

After cryoSPARC live pre-processing, 100 micrographs from each dataset were patch motion corrected and used to generate a denoiser model which was then used to denoise all micrographs within the dataset.

#### Hemagglutinin

For all HA datasets (0×, 0.25×, 1× SurfACT D21 H3N2 HA and 0×, 0.25× SurfACT CA H1N1 HA) denoised micrographs were blob-picked in cryoSPARC^77^ using a circular and elliptical blob. Particles were extracted at a box size of 384 pixels and Fourier cropped to 96 pixels at 4.41 Å/pixel. Particles were then subjected to two rounds of 2-D classification, where obvious HA classes were chosen to move forward. Two class *ab-initio* was then used as a particle pruning step as well as to generate an initial volume for further refinements.^77^ The selected particles underwent ANTIDOTE particle curation within RELION,^78,79^ then were exported back to cryoSPARC and subjected to non-uniform (NU) refinement before reextraction with a box size of 448 pixels at 0.735 Å/pixel. The final unbinned particle stack underwent reference-based motion correction (RBMC) before final NU refinements in both C_1_ and C_3_ symmetry (**Figure S2-6**).^80^

#### MoFeP

For all MoFeP datasets (0×, 0.25×, and 1× SurfACT) denoised micrographs were used for template-based particle picking in cryoSPARC using a template generated from a previously published MoFeP model (EMDB: 48384).^55^ Picked particles were first filtered based on confidence metric and extracted at a box size of 384 pixels with Fourier cropping to 96 pixels (4.41 Å/pixel). Particles were then subjected to two rounds of 2-D classification, where obvious MoFeP classes were chosen to move forward. *Ab-initio* model generation was used to generate an initial volume for subsequent NU refinement.^77,80^ Particles underwent ANTIDOTE particle curation within RELION, then were exported back to cryoSPARC and re-extracted with a box size of 384 pixels (0.735 Å/pixel) before NU refinement with C_1_ or C_2_ symmetry imposed.^78,79^ The C_1_ refined volume represented a “broken” incomplete MoFeP, while the C_2_ refined volume represented an “intact” complete MoFeP. The two refined volumes were used as initial volumes in a heterogenous refinement to sort for “intact” and “broken” particles. The “intact” MoFeP class was used as the final particle stack and underwent RBMC before final NU refinements in both C_1_ and C_2_ (**Figure S9-10**).^80^

#### Aldolase

For all aldolase datasets (0×, 0.25×, 0.5×, 1× SurfACT chameleon, 1× SurfACT gold-coated grid chameleon, 0×, 1× SurfACT Vitrobot, and 0.25× SurfACT manually-operated plunger), denoised micrographs were used for template-based particle picking in cryoSPARC using a template generated from a previously published aldolase study (EMDB: 21023).^70^ Picked particles were first filtered based on confidence metric and extracted at a box size of 256 pixels with Fourier cropping to 64 pixels (2.94 Å/pixel). Particles were then subjected to two rounds of 2-D classification, where obvious aldolase classes were chosen for downstream processing. Two class *ab-initio* was then used as a particle pruning step, as well as to generate an initial volume for further refinements.^77^ The selected particles underwent ANTIDOTE particle curation within RELION, then were exported back to cryoSPARC and subjected to (NU) refinement before re-extraction with a box size of 384 pixels (0.735 Å/pixel).^78,79^ The fully unbinned particles underwent NU refinement with C_1_ or D_2_ symmetry imposed. The C_1_ refined volume represented a “broken” protomer aldolase, while the D_2_ refined volume represented an “intact” aldolase with all four protomers. The two refined volumes were used as initial volumes in a heterogenous refinement to sort for “intact” and “broken” particles. The “intact” aldolase class was used as the final particle stack and underwent RBMC before final NU refinements in both C_1_ and D_2_ (**Figure S11-21**).^80^

### Model Building and Refinement

All HA, MoFeP, and aldolase models within this work were generated using a similar workflow. First, final volumes were subjected to the phenix.resolve cryo em density modification tool.^81^ Only half maps and sequence files were supplied, and default values were used. For each HA sample, ModelAngelo^82^ was employed to generate an initial structure using the same sequence as used for phenix.resolve cryo em. These initial models were fit into the resulting RESOLVE density and manually adjusted in COOT.^83^ For MoFeP structures, PDB: 7UT7 was fit into the RESOLVE density and manually adjusted in COOT. For aldolase maps, PDB: 6V20 was fit into the RESOLVE density and manually adjusted in COOT. Second, real-space refinements were performed in the RESOLVE densities using Phenix.^81^ For MoFeP, parameter files for the P-cluster (P^*N*^) and homocitrate-FeMoco ligands were used during refinement. Thirdly, each model from the resulting refinement was then treated with phenix.douse using the sharpened map from cryoSPARC to identify and add waters. Finally, the model was checked for accuracy in COOT, and a final real-space refinement was performed against the sharpened map from cryoSPARC (**Supplementary Table 2-3, 5-8**).

### CryoET Data Collection

Tilt series data collection was acquired at UCSD’s CryoEM Facility using a Titan Krios G4 (Thermo Fisher Scientific) operating at 300 kV equipped with a Selectris-X energy filter (Thermo Fisher Scientific). Target selection and data acquisition were performed by Tomo5 (Thermo Fisher Scientific). Data were acquired at a nominal magnification of 64,000× with a pixel size of 2.11 Å and a nominal defocus of *−*3.5 *µ*m. Tilt series collection was done in a dose-symmetric scheme with 3° tilt increment and angles ranging from *−*54° and +54°. The data was saved in the EER format with a total dose per tilt of ~2.68 e/Å^2^ and exposure time of 1.96s (**Supplementary Table 4, 9-10**).

### CryoET Data Processing

For all tomography datasets (0×, 0.25× SurfACT D21 H3N2 HA and 0×, 0.25×, 1× SurfACT chameleon-frozen aldolase), motion correction and CTF estimation were performed in WarpTool,^84^ prior to tilt-series alignment using patch-tracking in batch model in Etomo.^85^ The resulting alignment files from Etomo were imported into Warp using ts import alignment program, and the tomograms were reconstructed using WarpTools for template-matching. pyTOM-match-pick^86,87^ was used for template-matching and AMIRA (Thermo Fisher Scientific) was used to make manual masks and limit the extraction of candidates visually. Tomograms were denoised using Noise2Map incorporating odd/even half-maps of the tomogram, which were thresholded to further limit pyTOM’s particle extraction. Star files from pyTOM where then cleaned up in Cube (developed by Dimitry Tegunov) manually, and particles were extracted using the 2-D stack extraction in WarpTools and refined in RELION-5.^88^ To calculate the distance of particles from the AWIs, AMIRA was used to segment the tomograms to locate the top and bottom surfaces, and an in-house Python code was used to calculate the distances. A threshold of 100 Å from the mask was used to identify particles at the “top” AWI and “bottom” AWI, and remaining particles were designated “center” ice. Duplicate particles were also removed in the same code. Final particle positions and orientations were mapped back to the original tomogram using ArtiaX extension of ChimeraX.^89,90^ Aldolase subvolume averaging was done using C_1_ symmetry, but the HA trimers were refined using C_3_ symmetry since the preferred orientation was on the symmetry axis and the degradation was unaffected through averaging (**Figure S8**).

### Aldolase CryoET Particle Concentration Calculation

To estimate the concentration of aldolase in our tomograms we used a Python code to calculate the volume and particle count within a given mask used for data processing (see *CryoET Data Processing*). For the overall tomogram particle concentrations, the full ice mask and particle stack was used. For segmented tomogram particle concentrations, the code counts number of voxels within 100 Å of each ice surface, and the particles in those areas are assigned to the top and bottom AWI. The areas that are further than 100 Å from both surfaces are considered the center ice. We multiplied the number of voxels by 10.55 Å^3^ (the voxel size of the segmentation) to calculate vitreous ice volumes and the number of particles by 160 kDa to get the mass. The density was then converted to mg/mL and compared to applied sample concentration during grid preparation (**Figure S23**).

## Data Availability

The data that support this study are available from the corresponding authors upon request. CryoEM maps are available at the Electron Microscopy Data Bank (EMDB) with accession codes:

EMD-75842 (D21 H3N2 HA with 0× SurfACT, C_1_), EMD-75843 (D21 H3N2 HA with 0× SurfACT, C_3_), EMD-75844 (D21 H3N2 HA with 0.25× SurfACT, C_1_), EMD-75845 (D21 H3N2 HA with 0.25× SurfACT, C_3_), EMD-75846 (D21 H3N2 HA with 1× SurfACT, C_1_), EMD-75847 (D21 H3N2 HA with 1× SurfACT, C_3_), EMD-75848 (CA H1N1 HA with 0× Sur-fACT, C_1_), EMD-75849 (CA H1N1 HA with 0× SurfACT, C_3_), EMD-75851 (CA H1N1 HA with 0.25× SurfACT, C_1_), EMD-75853 (CA H1N1 HA with 0.25× SurfACT, C_3_), EMD-75854 (MoFeP with 0× SurfACT, C_1_), EMD-75855 (MoFeP with 0× SurfACT, C_2_), EMD-75856 (MoFeP with 0.25× Sur-fACT, C_1_), EMD-75857 (MoFeP with 0.25× SurfACT, C_2_), EMD-75858 (MoFeP with 1× SurfACT, C_1_), EMD-75859 (MoFeP with 1× SurfACT, C_2_), EMD-75860 (Aldolase with 0× SurfACT frozen on chameleon, C_1_), EMD-75861 (Aldolase with 0× SurfACT frozen on chameleon, D_2_), EMD-75862 (Aldolase with 0.25× SurfACT frozen on chameleon, C_1_), EMD-75863 (Aldolase with 0.25× SurfACT frozen on chameleon, D_2_), EMD-75864 (Aldolase with 0.5× SurfACT frozen on chameleon, C_1_), EMD-75865 (Aldolase with 0.5× SurfACT frozen on chameleon, D_2_), EMD-75866 (Aldolase with 1× Sur-fACT frozen on chameleon, C_1_), EMD-75867 (Aldolase with 1× SurfACT frozen on chameleon, D_2_), EMD-75868 (Aldolase with 1× SurfACT frozen on chameleon gold-coated grids, C_1_), EMD-75869 (Aldolase with 1× SurfACT frozen on chameleon gold-coated grids, D_2_), EMD-75870 (Aldolase with 0× Sur-fACT frozen on Vitrobot, C_1_), EMD-75871 (Aldolase with 0× SurfACT frozen on Vitrobot, D_2_), EMD-75873 (Aldolase with 1× SurfACT frozen on Vitrobot, C_1_), EMD-75874 (Aldolase with 1× SurfACT frozen on Vitrobot, D_2_), EMD-75878 (Aldolase with 0.25× SurfACT frozen on manual-plunge device, C_1_), EMD-75879 (Aldolase with 0.25× SurfACT frozen on manual-plunge device, D_2_). EMD-75901 (D21 H3N2 HA with 0× SurfACT, STA, C_3_), EMD-75902 (D21 H3N2 HA with 0.25× SurfACT, STA, C_3_), EMD-75903 (Aldolase with 0× SurfACT frozen on chameleon, STA, C_1_), EMD-75904 (Aldolase with 0.25× SurfACT frozen on chameleon, STA, C_1_), EMD-75905 (Aldolase with 0.25× SurfACT at top AWI, STA, C_1_), EMD-75906 (Aldolase with 0.25× SurfACT in center ice, STA, C_1_), EMD-75907 (Aldolase with 0.25× SurfACT at bottom AWI, STA, C_1_), EMD-75908 (Aldolase with 1× SurfACT frozen on chameleon, STA, C_1_).

Corresponding structural models have been deposited in the Protein Data Bank (PDB) with accession codes:

11MS (D21 H3N2 HA with 0.25× SurfACT, C_1_), 11MT (D21 H3N2 HA with 0.25× SurfACT, C_3_), 11MU (D21 H3N2 HA with 1× SurfACT, C_1_), 11MV (D21 H3N2 HA with 1× SurfACT, C_3_), 11MX (CA H1N1 HA with 0.25× SurfACT, C_1_), 11MZ (CA H1N1 HA with 0.25× SurfACT, C_3_), 11NA (MoFeP with 0× SurfACT, C_1_), 11NB (MoFeP with 0× Sur-fACT, C_2_), 11NC (MoFeP with 0.25× SurfACT, C_1_), 11ND (MoFeP with 0.25× SurfACT, C_2_), 11NE (MoFeP with 1× SurfACT, C_1_), 11NF (MoFeP with 1× SurfACT, C_2_), 11NH (Aldolase with 0× SurfACT frozen on chameleon, C_1_), 11NI (Aldolase with 0× SurfACT frozen on chameleon, D_2_), 11NJ (Aldolase with 0.25× SurfACT frozen on chameleon, C_1_), 11NK (Aldolase with 0.25× SurfACT frozen on chameleon, D_2_), 11NL (Aldolase with 0.5× SurfACT frozen on chameleon, C_1_), 11NM (Aldolase with 0.5× SurfACT frozen on chameleon, D_2_), 11NN (Aldolase with 1× SurfACT frozen on chameleon, C_1_), 11NO (Aldolase with 1× SurfACT frozen on chameleon, D_2_), 11NP (Aldolase with 1× SurfACT frozen on chameleon gold-coated grids, C_1_), 11NR (Aldolase with 1× SurfACT frozen on chameleon gold-coated grids, D_2_), 11NT (Aldolase with 0× SurfACT frozen on Vitrobot, C_1_), 11NU (Aldolase with 0× SurfACT frozen on Vitrobot, D_2_), 11NW (Aldolase with 1× SurfACT frozen on Vitrobot, C_1_), 11NX (Aldolase with 1× SurfACT frozen on Vitrobot, D_2_), 11OB (Aldolase with 0.25× SurfACT frozen on manual-plunge device, C_1_), 11OC (Aldolase with 0.25× SurfACT frozen on manual-plunge device, D_2_). All other data are available in the main text or the Supplementary Materials.

## Acknowledgements

We are grateful to Dr. Kevin Corbett for providing valuable feedback on this manuscript, and the entirety of the Herzik lab for facilitating insightful discussions. The authors acknowledge the cryoEM facility at UC San Diego, along with the scientific and technical assistance of the facility director, Dr. Mariusz Matyszewski. We also thank Brendan Dennis, Kevin Smith, and the UCSD Physics Computing Facility for their insights and support. We would also like to thank JC Ducom and Lisa Dong from Scripps Research High-Performance Computing for their support. We are grateful to Dr. Julianna Han, Alesandra Rodriguez, and Dr. Gabriel Ozorowski from the Ward lab for providing the California 2009 H1N1 and Darwin 2021 H3N2 HA samples. Molecular graphics and analyses were performed with UCSF ChimeraX, developed by the Resource for Biocomputing, Visualization, and Informatics at the University of California, San Francisco (UCSF), with support from National Institutes of Health (NIH) grant R01-GM129325 and the Office of Cyber Infrastructure and Computational Biology, National Institute of Allergy and Infectious Diseases. This work was supported by the NIH grant R35-GM138206, Searle Scholars program, Cottrell Scholars program, and George W. and Carol A. Lattimer Faculty Research Fellowship (M.A.H.) and NIH grant GM154216-02 (D.A.G). The chameleon used in this protocol was obtained via NIH 1S10OD032471 (M.A.H.). S.E.E. and M.J.B were supported by the NIH Molecular Biophysics Training Program at UCSD (T32-GM139795). B.D.C. was supported by a Goeddel Family Technology Sandbox Fellowship. S.M.N. was supported by the NIH Interfaces Graduate Training Program at UCSD (T32-EB009380).

## Contributions

S.E.E. and B.D.C. conceived the project, designed experiments, performed cryoEM experiments and analysis, and co-wrote the manuscript. H.R. performed cryoET data processing and analysis. S.M.N. performed protein isolation/characterization for MoFeP. I.C.K. performed Vitrobot cryoEM experiments. Y.L. and M.J.B. developed the Sur-fACT formulation. B.D.C., H.T.R., P.D., and D.J. surveyed the EMDB. D.A.G. provided guidance in cryoET experimental design and analysis, manuscript writing, and project management. M.A.H. conceived and directed the project, designed and oversaw cryoEM experiments, and co-wrote the manuscript. All authors participated in the editing of the manuscript.

## Ethics Declarations

The authors declare no competing interests.

