## Supplemental Files for "A Surfactant Cocktail Overcomes Air-Water Interface Artifacts in Single-Particle CryoEM"

| Surfactant | Chemical Structure | Critical Micelle Concentration (w/v) | 1× SurfACT (w/v) | 0.5× SurfACT (w/v) | 0.25× SurfACT (w/v) |
|---|---|---|---|---|---|
| FOM *(small, non-ionic detergent)* | 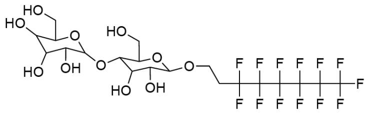 | 0.048% | 0.0048% | 0.0024% | 0.0012% |
| Brij-35 *(large, non-ionic detergent)* | 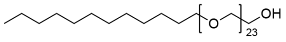 | 0.6% | 0.06% | 0.03% | 0.015% |
| CHAPSO *(zwitterionic detergent)* | 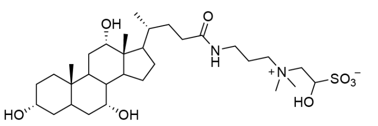 | 0.5% | 0.05% | 0.025% | 0.0125% |
| A8-35 *(amphiphilic polymer)* | 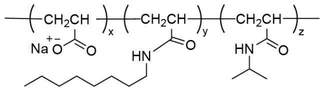 | 2.5% | 0.25% | 0.125% | 0.0625% |

**Figure S1: Chemical structures and properties of SurfACT components**
SurfACT composition, including fluorinated octyl maltoside (FOM), Brij-35, 3-[(3-cholamidopropyl)-dimethylammonio]-2-hydroxy-1-propanesulfonate (CHAPSO), and amphipol 8-35 (A8-35) based on our survey of surfactant usage for preferred orientation in the EMDB (see **Figure 1** and **Methods**). For each component, the physicochemical property, chemical structure, critical micelle concentration (CMC; protein-solubilizing concentration for A8-35), SurfACT formulation working concentration (FWC). *For A8-35, we used the protein-solubilizing concentration (∼5× protein concentration) based on an average protein concentration for cryoEM of ∼5 mg/ml. 0.5× and 0.25× surfACT concentrations are calculated relative to the FWC.

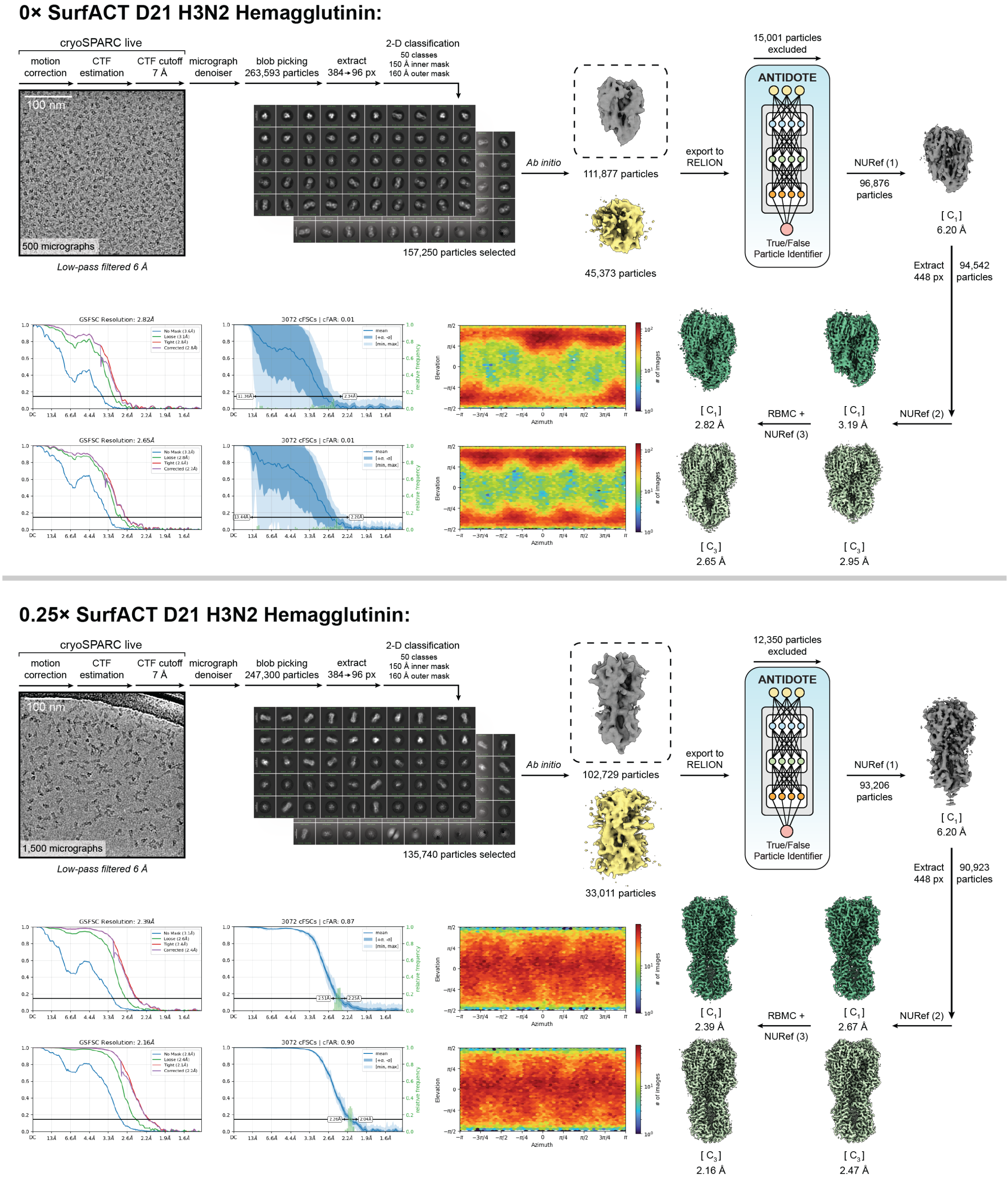


**Non-uniform refinement (NURef) 1**: minimize over per-particle scale, initialize noise model from images. **NURef 2**: minimize over per-particle scale, initialize noise model from images, optimize per-particle defocus, optimize CTF parameters. **NURef 3**: no real-space window, minimize over per-particle scale, initialize noise model from images, optimize per-particle defocus, optimize CTF parameters. **Reference-Based Motion Correction (RBMC):** override EER number of fractions (80).

**Figure S2: CryoEM processing of D21 H3N2 hemagglutinin prepared with 0× and 0.25× SurfACT using a SPT Labtech chameleon** Single particle cryoEM workflow for D21 H3N2 hemagglutinin prepared with 0× and 0.25× SurfACT using a SPT Labtech chameleon. Representative aligned micrographs, 2-D class averages, and 3-D sorting and refinement steps in cryoSPARC and RELION are detailed.[1,2] ANTIDOTE[3] particle curation within RELION prior to final 3-D sorting and refinement in cryoSPARC is shown. Final $C_1$ and $C_3$ refinements, elevation plots, and 3DFSC curves are shown. Key parameters for non-uniform refinement (NUref)[4] and reference-based motion correction (RBMC) are shown.

### 1× SurfACT D21 H3N2 Hemagglutinin:

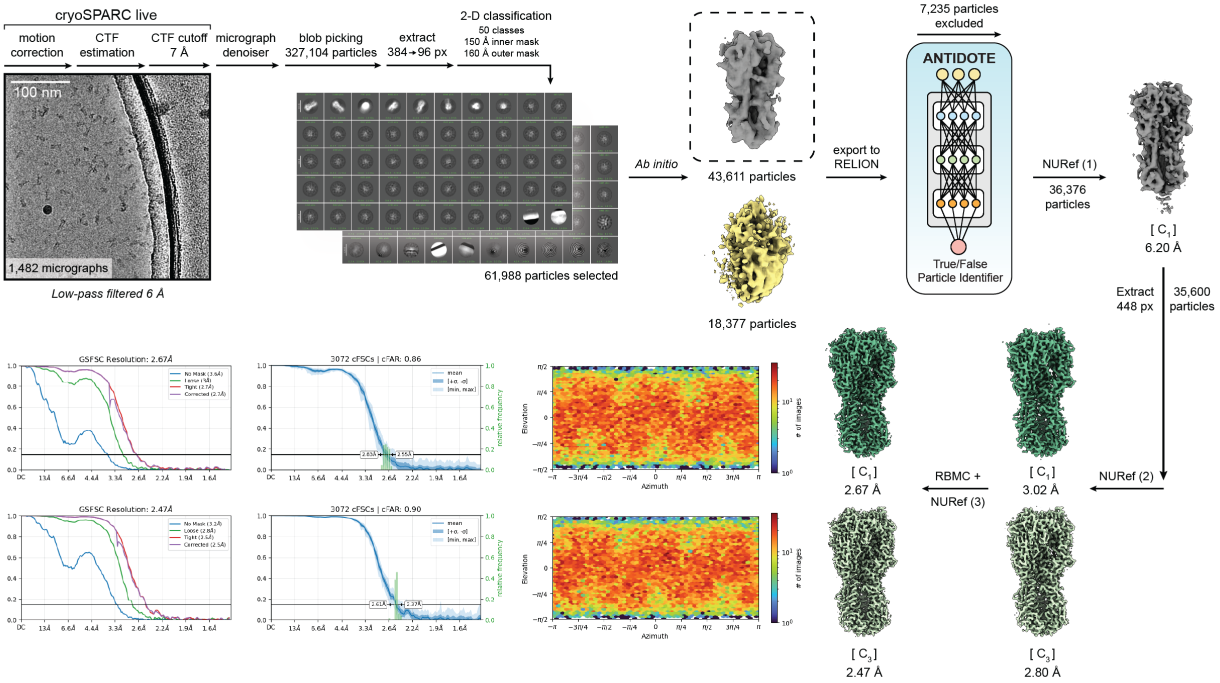


**Non-uniform refinement (NURef) 1**: minimize over per-particle scale, initialize noise model from images. **NURef 2**: minimize over per-particle scale, initialize noise model from images, optimize per-particle defocus, optimize CTF parameters. **NURef 3**: no real-space window, minimize over per-particle scale, initialize noise model from images, optimize per-particle defocus, optimize CTF parameters. **Reference-Based Motion Correction (RBMC):** override EER number of fractions (80).

**Figure S3: CryoEM processing of D21 H3N2 hemagglutinin prepared with 1× SurfACT using a SPT Labtech chameleon**
Single particle cryoEM workflow for D21 H3N2 hemagglutinin with 1× SurfACT prepared using a SPT Labtech chameleon. Representative aligned micrograph, 2-D class averages, and 3-D sorting and refinement steps in cryoSPARC and RELION are detailed.[1,2] ANTIDOTE[3] particle curation within RELION prior to final 3-D sorting and refinement in cryoSPARC is shown. Final $C_1$ and $C_3$ refinements, elevation plots, and 3DFSC curves are shown. Key parameters for non-uniform refinement (NUref)[4] and reference-based motion correction (RBMC) are shown.

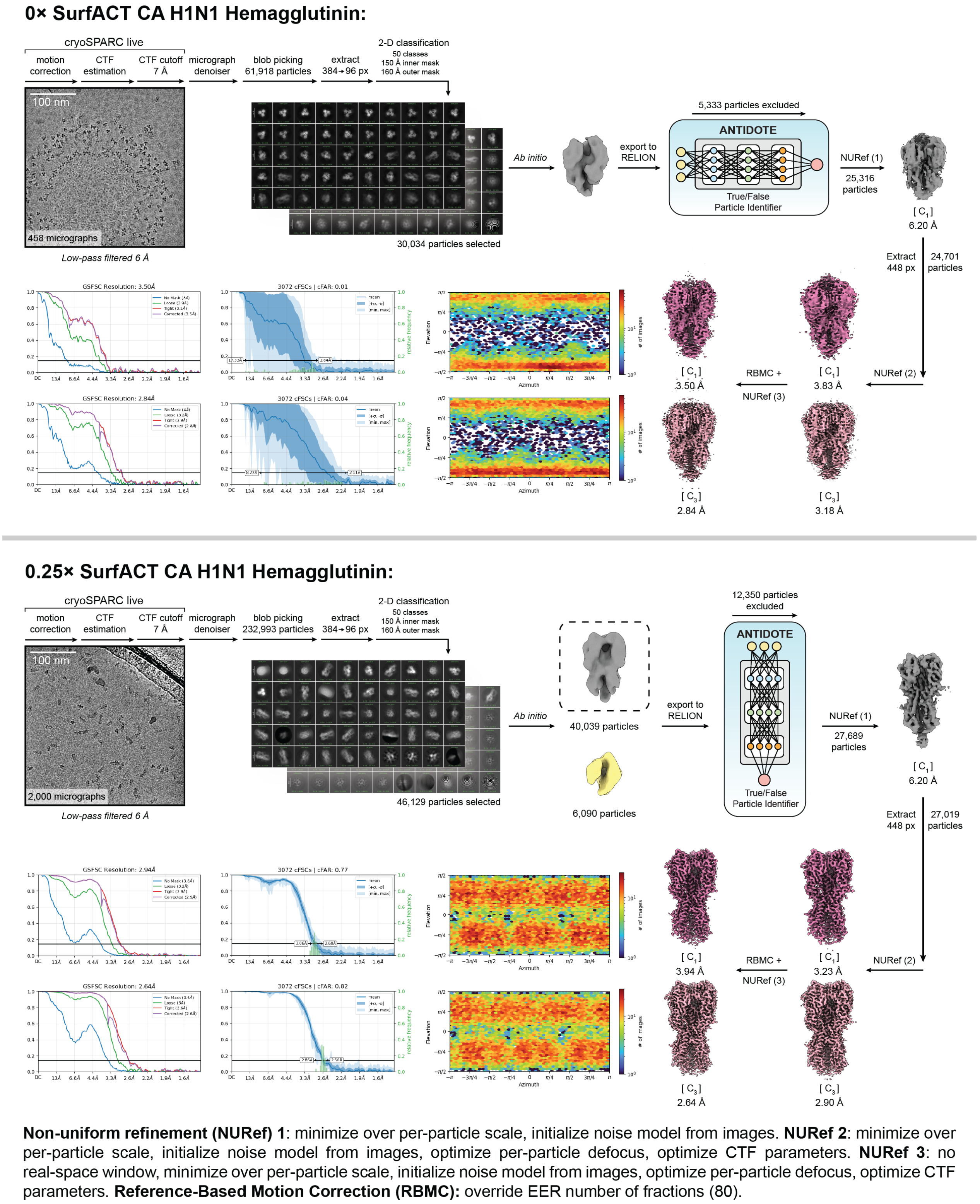


**Figure S4: CryoEM processing of CA H1N1 hemagglutinin prepared with 0× and 0.25× SurfACT using a SPT Labtech chameleon** Single particle cryoEM workflow for CA H1N1 hemagglutinin prepared with 0× and 0.25× SurfACT using a SPT Labtech chameleon. Representative aligned micrographs, 2-D class averages, and 3-D sorting and refinement steps in cryoSPARC and RELION are detailed.[1,2] ANTIDOTE[3] particle curation within RELION prior to final 3-D sorting and refinement in cryoSPARC is shown. Final $C_1$ and $C_3$ refinements, elevation plots, and 3DFSC curves are shown. Key parameters for non-uniform refinement (NUref)[4] and reference-based motion correction (RBMC) are shown.

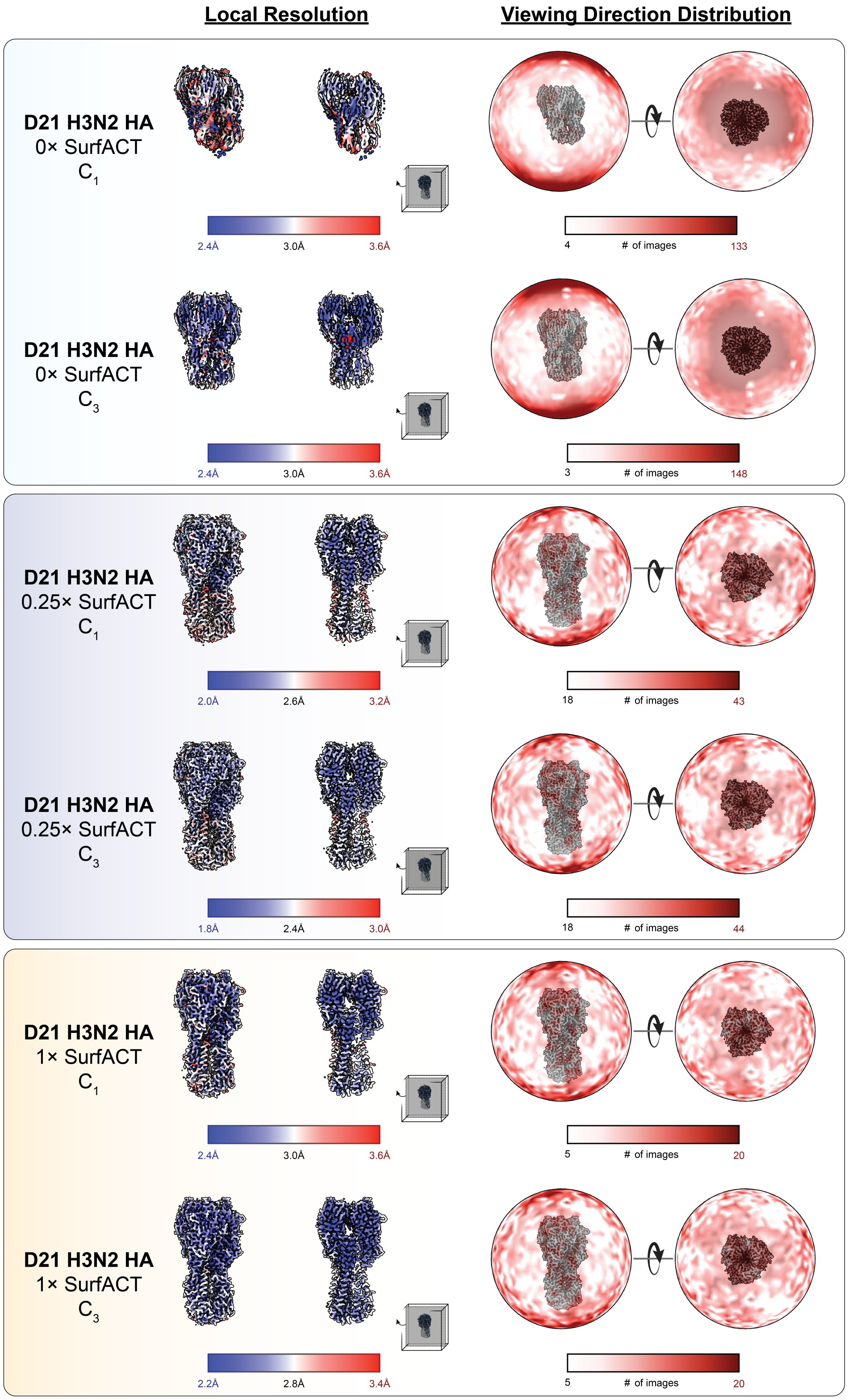


**Figure S5: Local Resolution and signal distribution for D21 H3N2 hemagglutinin prepared with 0×, 0.25×, and 1× SurfACT using a SPT Labtech chameleon**

EM density analysis for single particle cryoEM $C_1$ and $C_3$ reconstructions of D21 H3N2 hemagglutinin with 0×, 0.25×, and 1× SurfACT prepared using a SPT Labtech chameleon. Full volume and central slice of volume (*left*) are colored by local resolution blue(low)-white-red(high). Signal distribution (*right*) is shown as a heat map from white (low) to red (high).

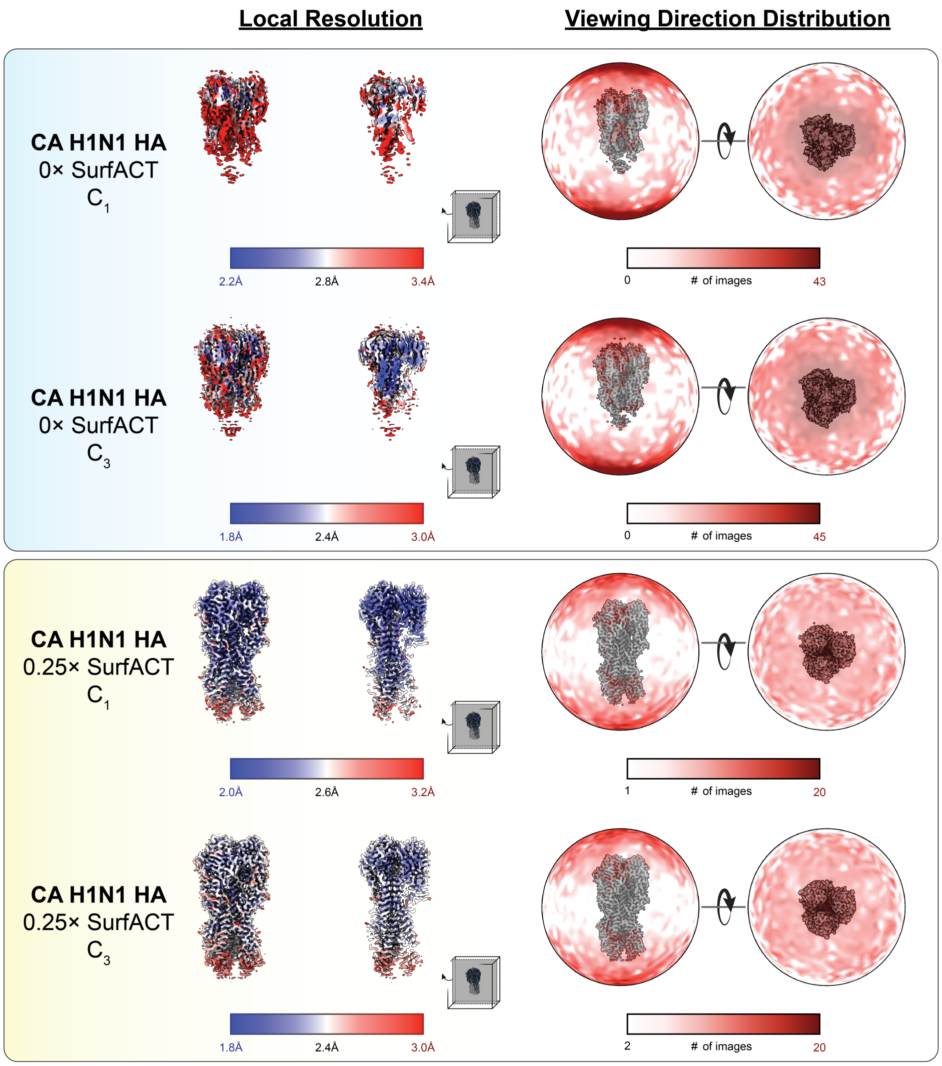


**Figure S6: Local Resolution and signal distribution for CA H1N1 hemagglutinin prepared with 0× and 0.25× SurfACT using a SPT Labtech chameleon**

EM density analysis for single particle cryoEM $C_1$ and $C_3$ reconstructions of CA H1N1 hemagglutinin with 0× and 0.25× SurfACT prepared using a SPT Labtech chameleon. Full volume and central slice of volume (*left*) are colored by local resolution blue(low)-white-red(high). Signal distribution (*right*) is shown as a heat map from white (low) to red (high).

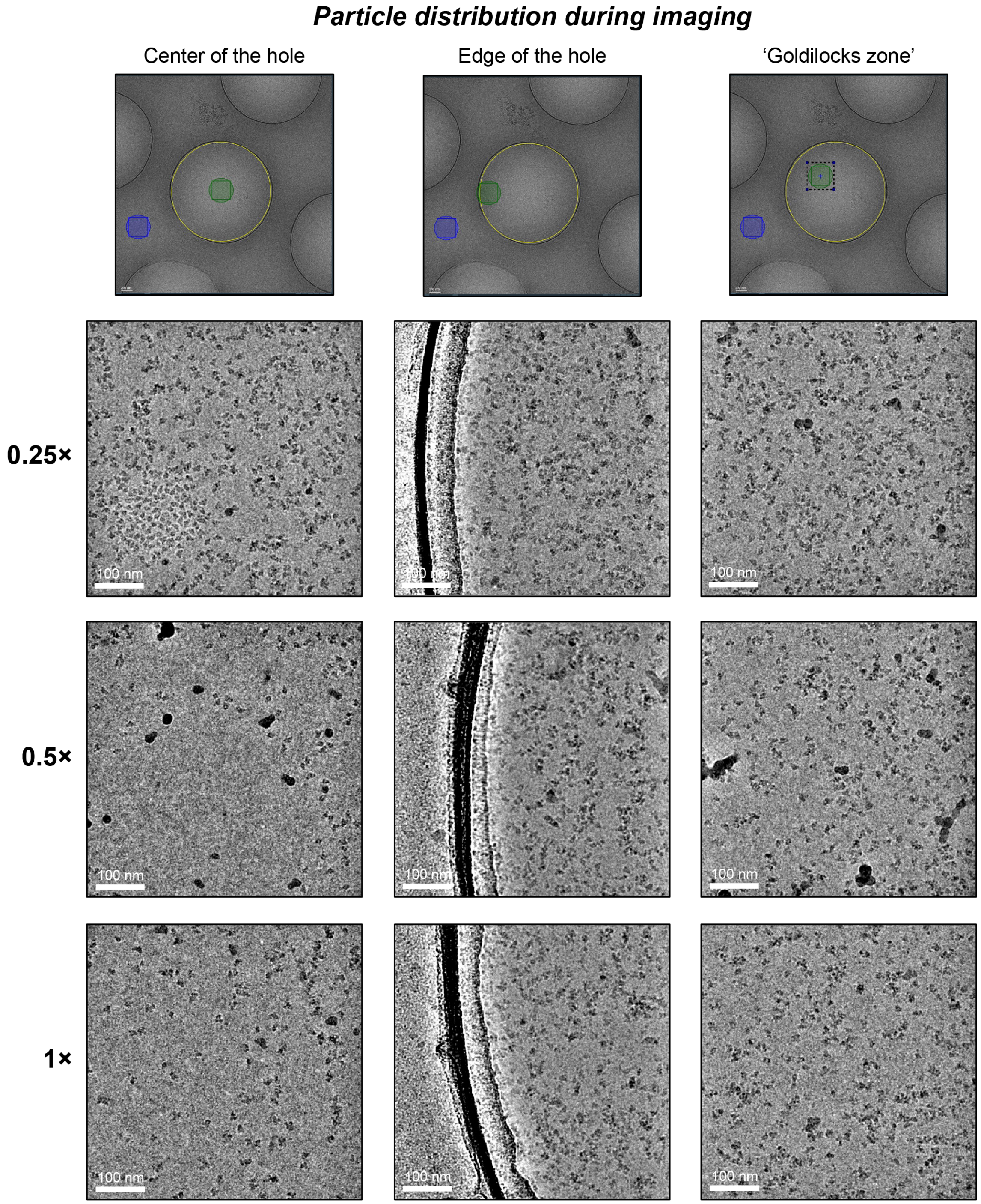


**Figure S7: Influence of varying SurfACT concentrations on aldolase particle density and distribution across cryoEM imaging regions using a SPT Labtech chameleon**

Micrographs of rabbit muscle aldolase (12 mg/ml initial concentration) frozen using a SPT Labtech chameleon with varying SurfACT concentrations (0.25×, 0.5×, 1×). With 0.25× SurfACT, particles are still present in the center of the grid hole, allowing for imaging in the thinnest ice region. With 0.5× and 1× SurfACT, particles are sparse in the center of the hole and locate more to the edges. By imaging in between the center and edge of the hole (*right*), we balance particle density and ice thickness for ideal data collection.

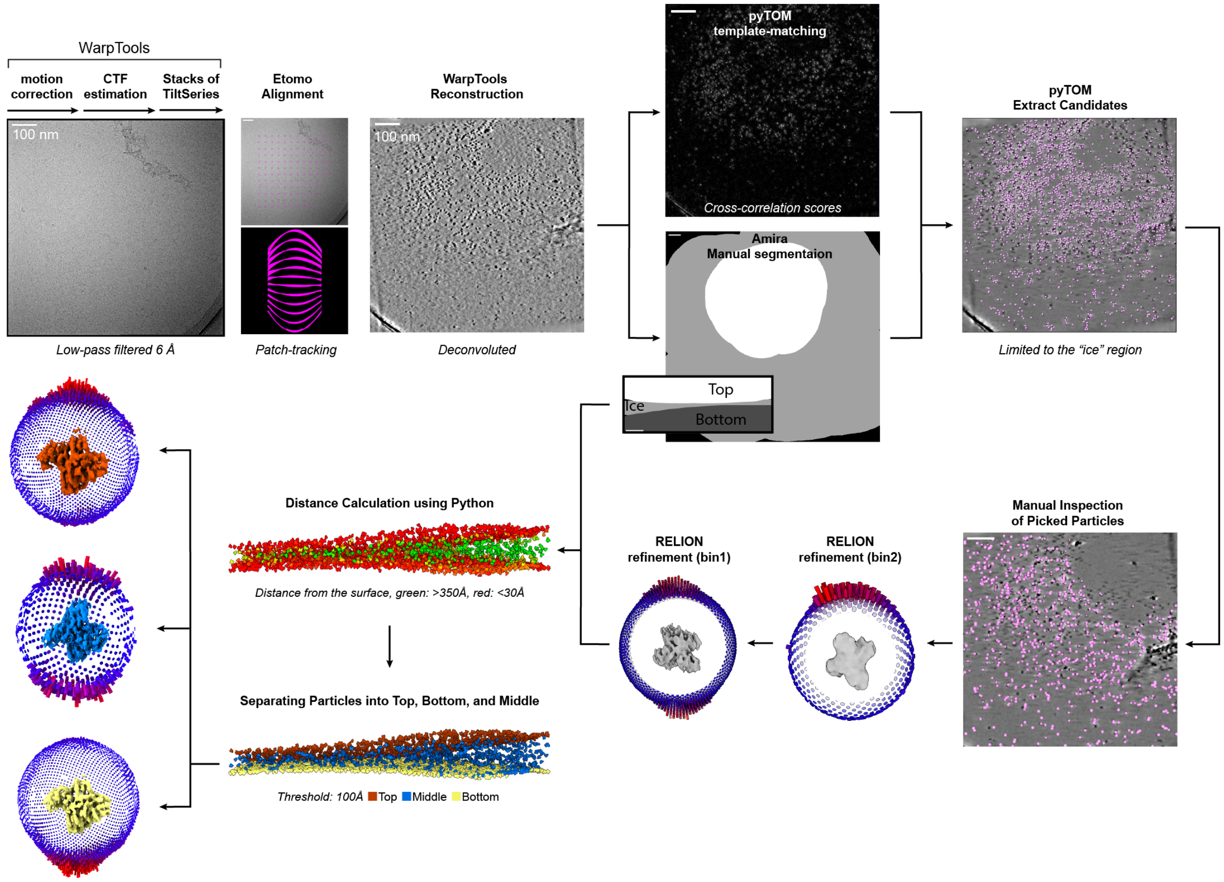


**Figure S8: CryoET data processing workflow for D21 H3N2 hemagglutinin (0× and 0.25× SurfACT) and aldolase (0×, 0.25×, and 1× SurfACT) datasets prepared using the chameleon**

Subtomogram averaging cryoET workflow for hemagglutinin and aldolase prepared using a chameleon. Etomo was used for tilt-series alignment,[5] followed by tomogram reconstruction using WarpTools.[6] Tomograms were picked using pyTOM paired with Amira to manually mask the vitreous ice.[7,8] Manually pruned particles were refined in RELION.[9] A similar workflow was used for all cryoET datasets (aldolase with 0×, 0.25×, and 1× SurfACT and HA with 0× and 0.25× SurfACT) with exception of the final top, middle, and bottom segmentation and separate reconstructions employed for 0.25× SurfACT aldolase.

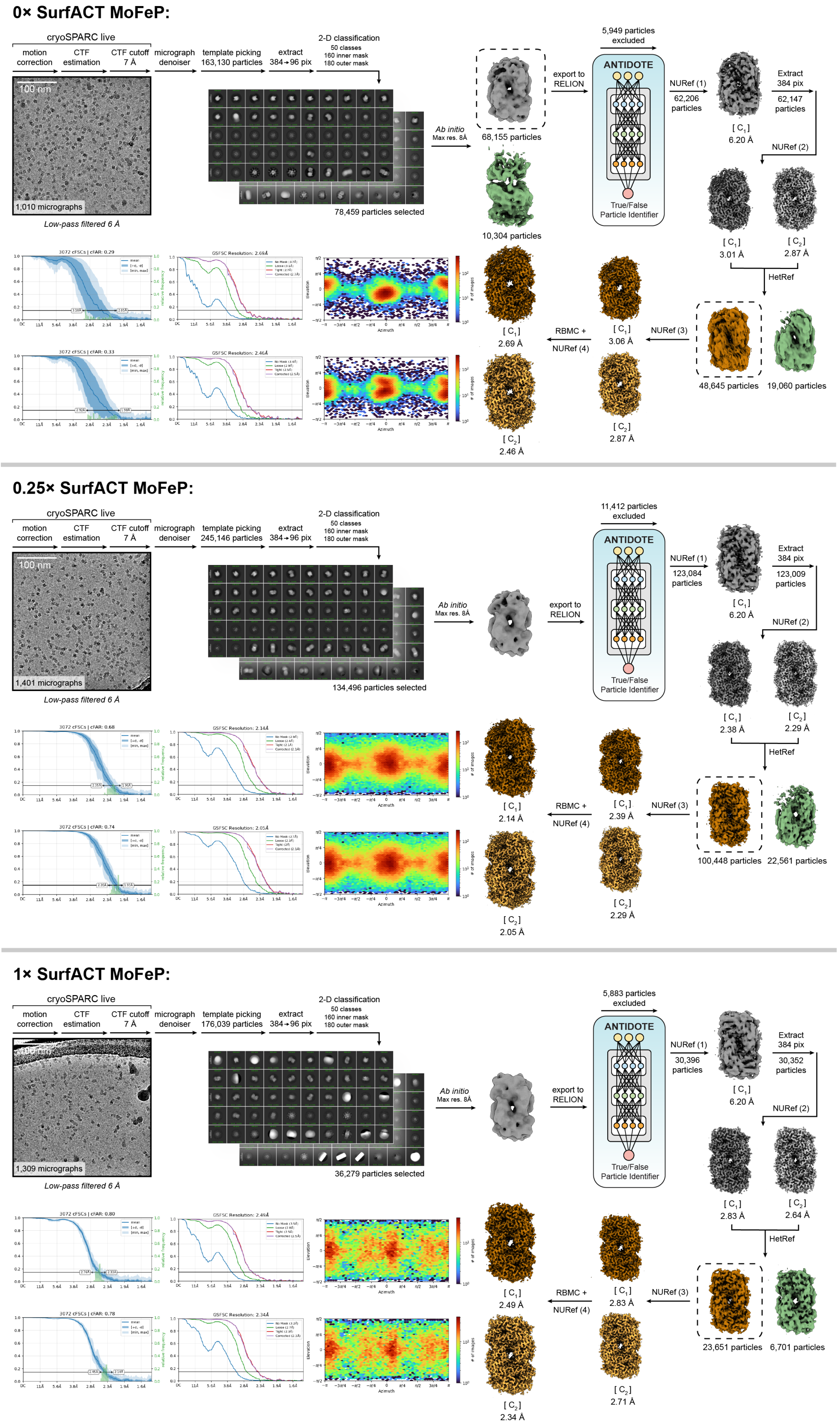


**Non-uniform refinement (NURef) 1**: minimize over per-particle scale, initialize noise model from images. **NURef 2-3**: minimize over per-particle scale, initialize noise model from images, optimize per-particle defocus, optimize CTF parameters. **NURef 4**: no real-space window, minimize over per-particle scale, initialize noise model from images, optimize per-particle defocus, optimize CTF parameters. **Heterogeneous refinement (HetRef)**: batch size per class 2000. **Reference-Based Motion Correction (RBMC):** override EER number of fractions (80).

**Figure S9: CryoEM processing of *Azotobacter vinellandii* MoFeP with 0× ,0.25×, and 1× SurfACT prepared using a SPT Labtech chameleon under anaerobic conditions**

Single particle cryoEM workflow for *Azotobacter vinellandii* MoFeP with 0×, 0.25×, and 1× SurfACT prepared using a chameleon under anaerobic conditions. Representative aligned micrographs, 2-D class averages, and 3-D sorting and refinement steps in cryoSPARC and RELION are detailed.[1,2] ANTIDOTE[3] particle curation within RELION prior to final 3-D sorting from heterogenous refinement (HetRef) in cryoSPARC with broken ($C_1$) and intact ($C_2$) volumes supplied. Final $C_1$ and $C_2$ refinements, elevation plots, and 3DFSC curves are shown. Key parameters for non-uniform refinement (NUref)[4] and reference-based motion correction (RBMC) are shown.

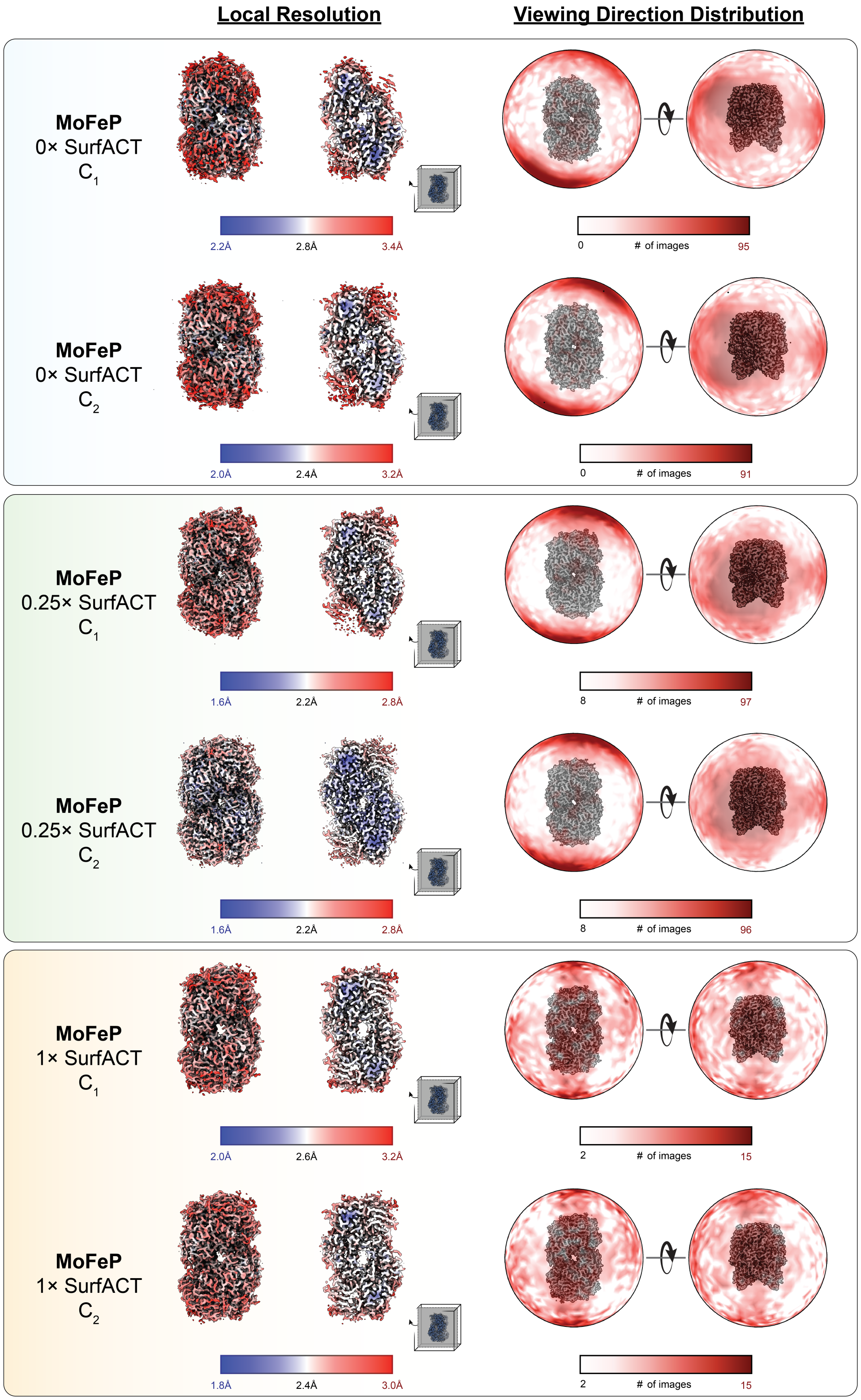


**Figure S10: Local Resolution and signal distribution for *Azotobacter vinellandii* MoFeP with 0×, 0.25×, and 1× SurfACT prepared using a SPT Labtech chameleon under anaerobic conditions**
EM density analysis for single particle cryoEM $C_1$ and $C_2$ reconstructions of Azotobacter vinellandii MoFeP with 0×, 0.25×, and 1× SurfACT prepared using a SPT Labtech chameleon under anaerobic conditions. Full volume and central slice of volume (*left*) are colored by local resolution blue(low)-white-red(high). Signal distribution (*right*) is shown as a heat map from white (low) to red (high).

### Aldolase with 0× SurfACT Frozen on the chameleon:

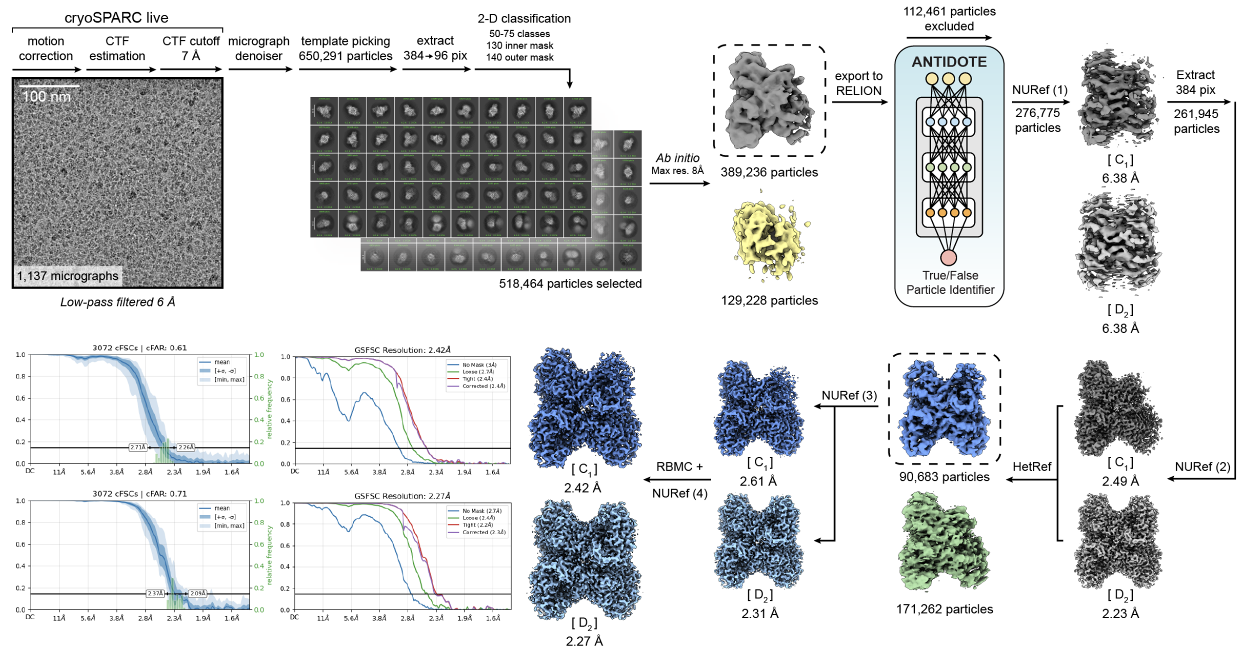


### Aldolase with 0.25× SurfACT Frozen on the chameleon:

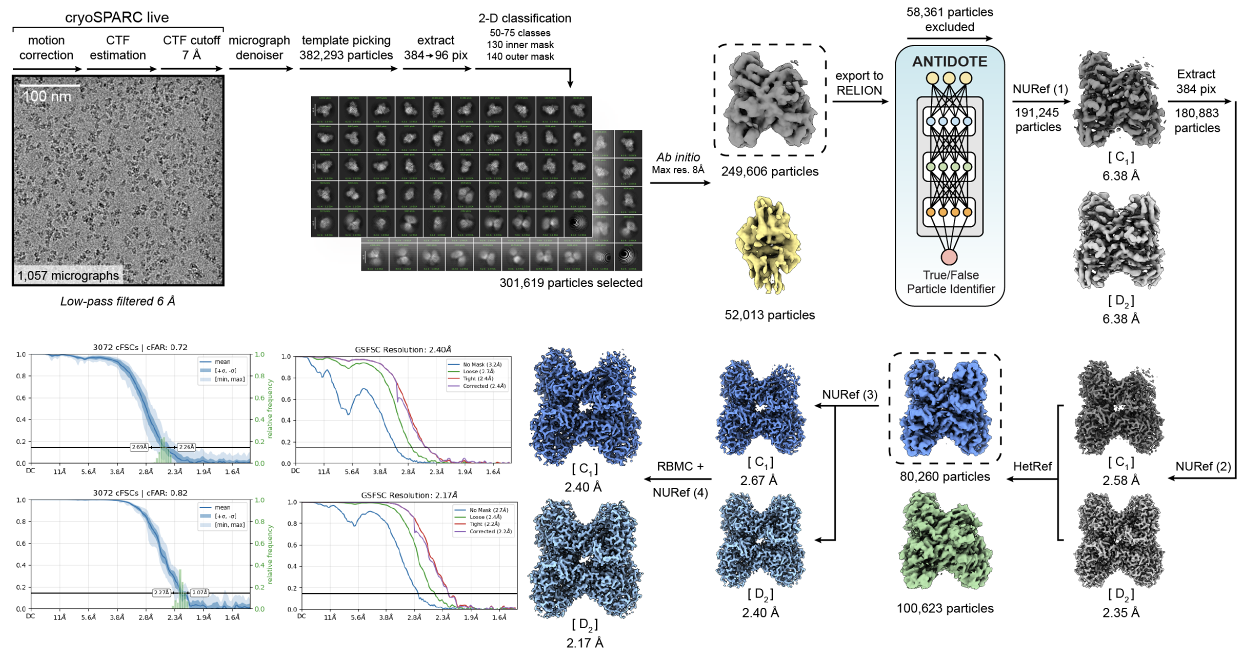


**Non-uniform refinement (NURef) 1**: minimize over per-particle scale, initialize noise model from images. **NURef 2-3**: minimize over per-particle scale, initialize noise model from images, optimize per-particle defocus, optimize CTF parameters. **NURef 4**: no real-space window, minimize over per-particle scale, initialize noise model from images, optimize per-particle defocus, optimize CTF parameters. **Heterogeneous refinement (HetRef)**: batch size per class 2000. **Reference-Based Motion Correction (RBMC):** override EER number of fractions (80).

**Figure S11: CryoEM processing of rabbit muscle aldolase prepared with 0× and 0.25× SurfACT using a SPT Labtech chameleon** Single particle cryoEM workflow for aldolase with 0× and 0.25× SurfACT prepared using a SPT Labtech chameleon. Representative aligned micrographs, 2-D class averages, and 3-D sorting and refinement steps in cryoSPARC and RELION are detailed.[1,2] ANTIDOTE[3] particle curation within RELION prior to final 3-D sorting from heterogenous refinement (HetRef) in cryoSPARC with broken ($C_1$) and intact ($D_2$) volumes supplied. Final $C_1$ and $D_2$ refinements, elevation plots, and 3DFSC curves are shown. Key parameters for non-uniform refinement (NUref)[4] and reference-based motion correction (RBMC) are shown.

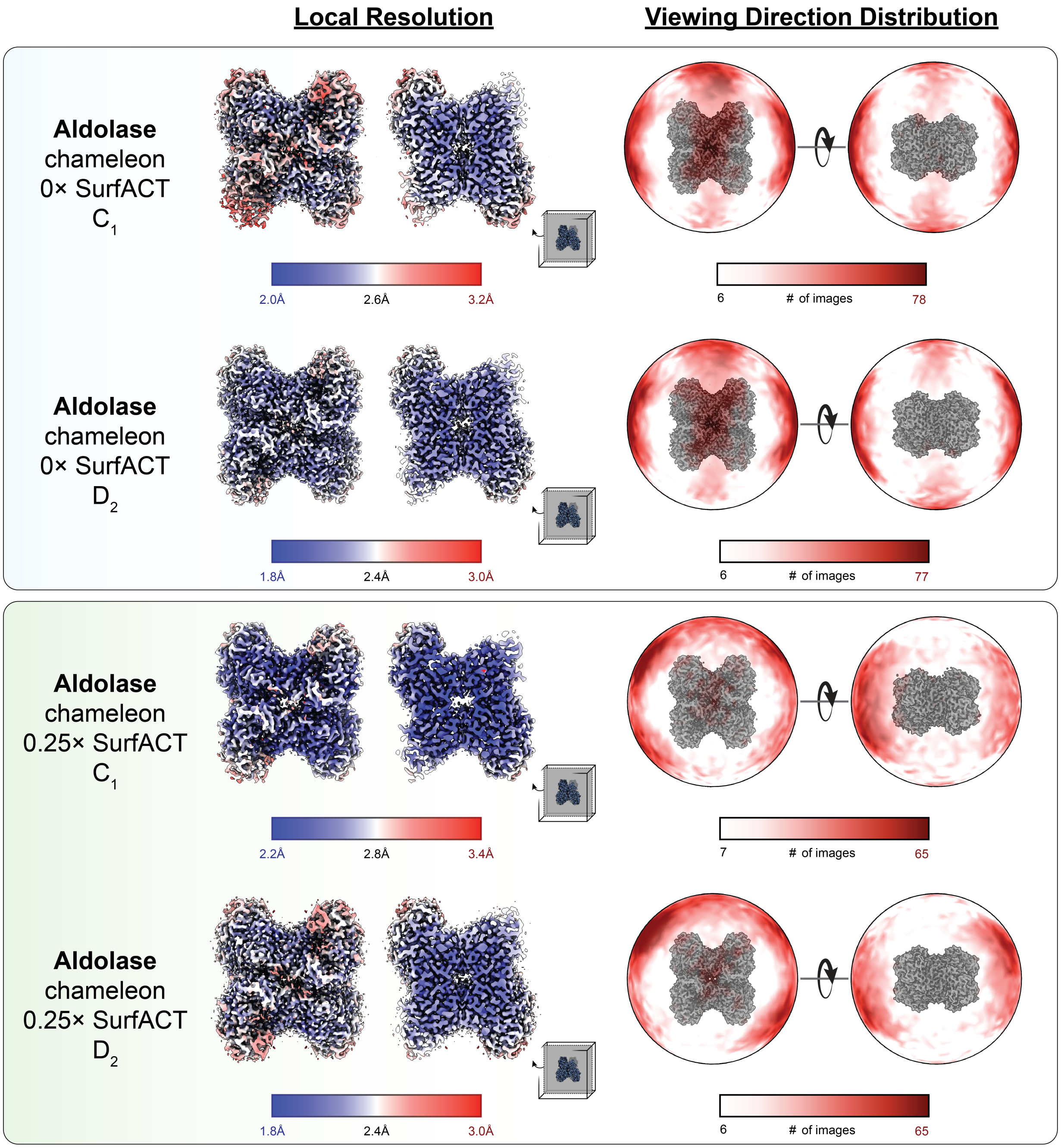


**Figure S12: Local Resolution and signal distribution for rabbit muscle aldolase prepared with 0× and 0.25× SurfACT using a SPT Labtech chameleon**

EM density analysis for single particle cryoEM $C_1$ and $D_2$ reconstructions of rabbit muscle aldolase with 0× and 0.25× SurfACT prepared using a SPT Labtech chameleon. Full volume and central slice of volume (*left*) are colored by local resolution blue (low)-white-red (high). Signal distribution (*right*) is shown as a heat map from white (low) to red (high).

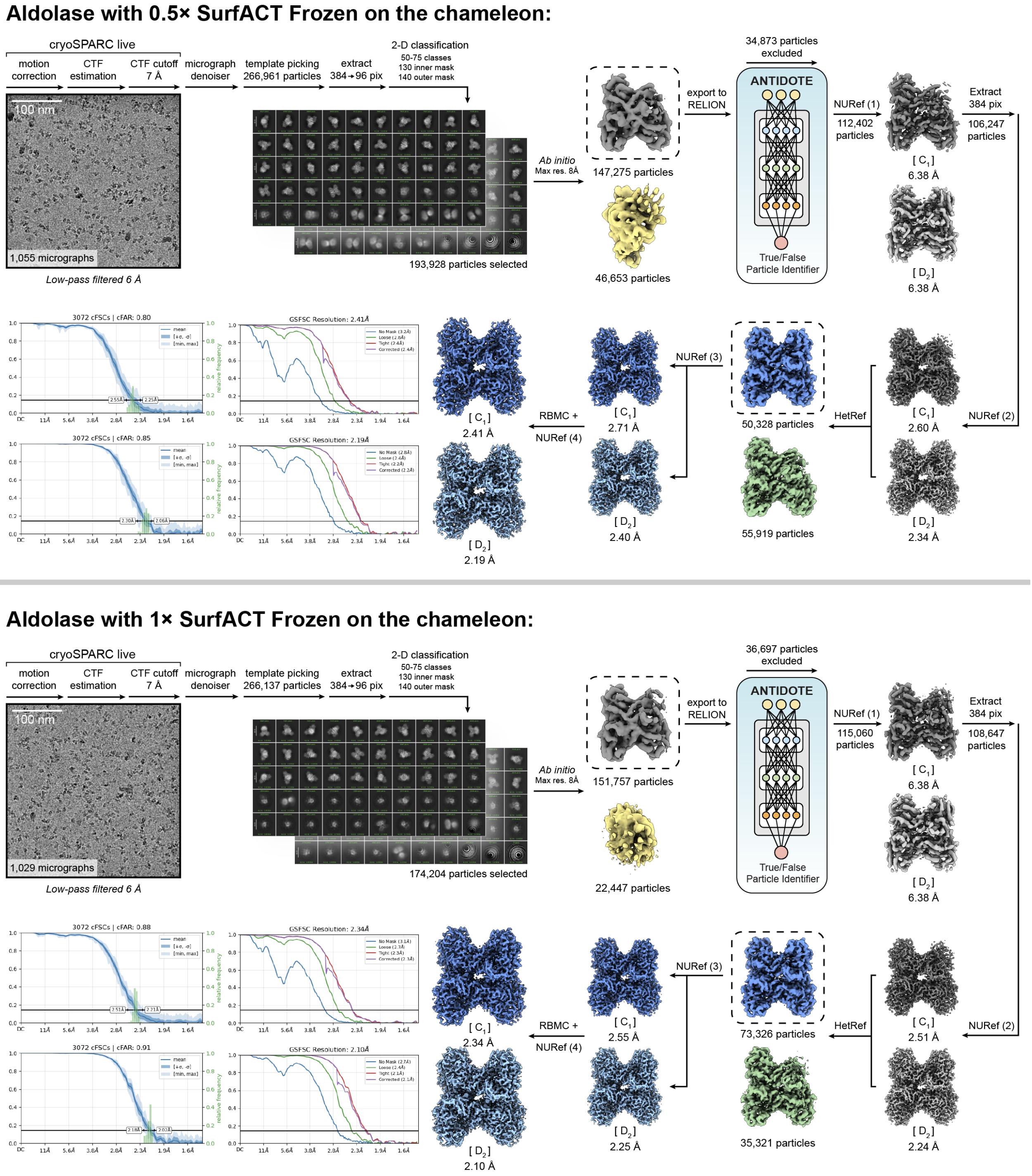


**Non-uniform refinement (NURef) 1**: minimize over per-particle scale, initialize noise model from images. **NURef 2-3**: minimize over per-particle scale, initialize noise model from images, optimize per-particle defocus, optimize CTF parameters. **NURef 4**: no real-space window, minimize over per-particle scale, initialize noise model from images, optimize per-particle defocus, optimize CTF parameters. **Heterogeneous refinement (HetRef)**: batch size per class 2000. **Reference-Based Motion Correction (RBMC):** override EER number of fractions (80).

**Figure S13: CryoEM processing of rabbit muscle aldolase prepared with 0.5× and 1× SurfACT using a SPT Labtech chameleon**
Single particle cryoEM workflow for aldolase with 0.5× and 1× SurfACT prepared using a SPT Labtech chameleon. Representative aligned micrographs, 2-D class averages, and 3-D sorting and refinement steps in cryoSPARC and RELION are detailed.[1,2] ANTIDOTE[3] particle curation within RELION prior to final 3-D sorting from heterogenous refinement (HetRef) in cryoSPARC with broken ($C_1$) and intact ($D_2$) volumes supplied. Final C1 and D2 refinements, elevation plots, and 3DFSC curves are shown. Key parameters for non-uniform refinement (NUref)[4] and reference-based motion correction (RBMC) are shown.

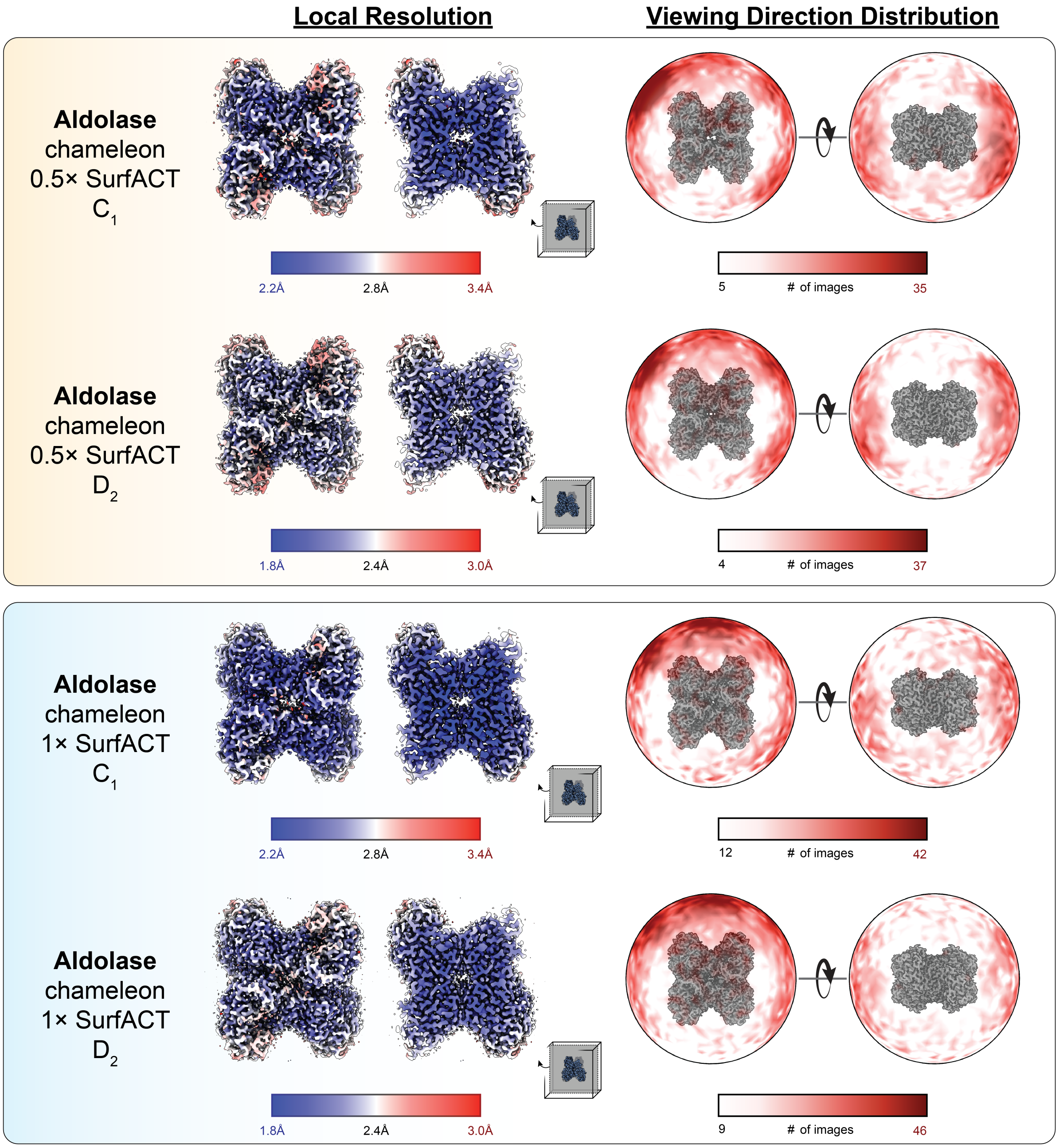


**Figure S14: Local Resolution and signal distribution for rabbit muscle aldolase prepared with 0.5× and 1× SurfACT using a SPT Labtech chameleon**
EM density analysis for single particle cryoEM $C_1$ and $D_2$ reconstructions of rabbit muscle aldolase with 0.5× and 1× SurfACT prepared using a SPT Labtech chameleon. Full volume and central slice of volume (*left*) are colored by local resolution blue (low)-white-red (high). Signal distribution (*right*) is shown as a heat map from white (low) to red (high).

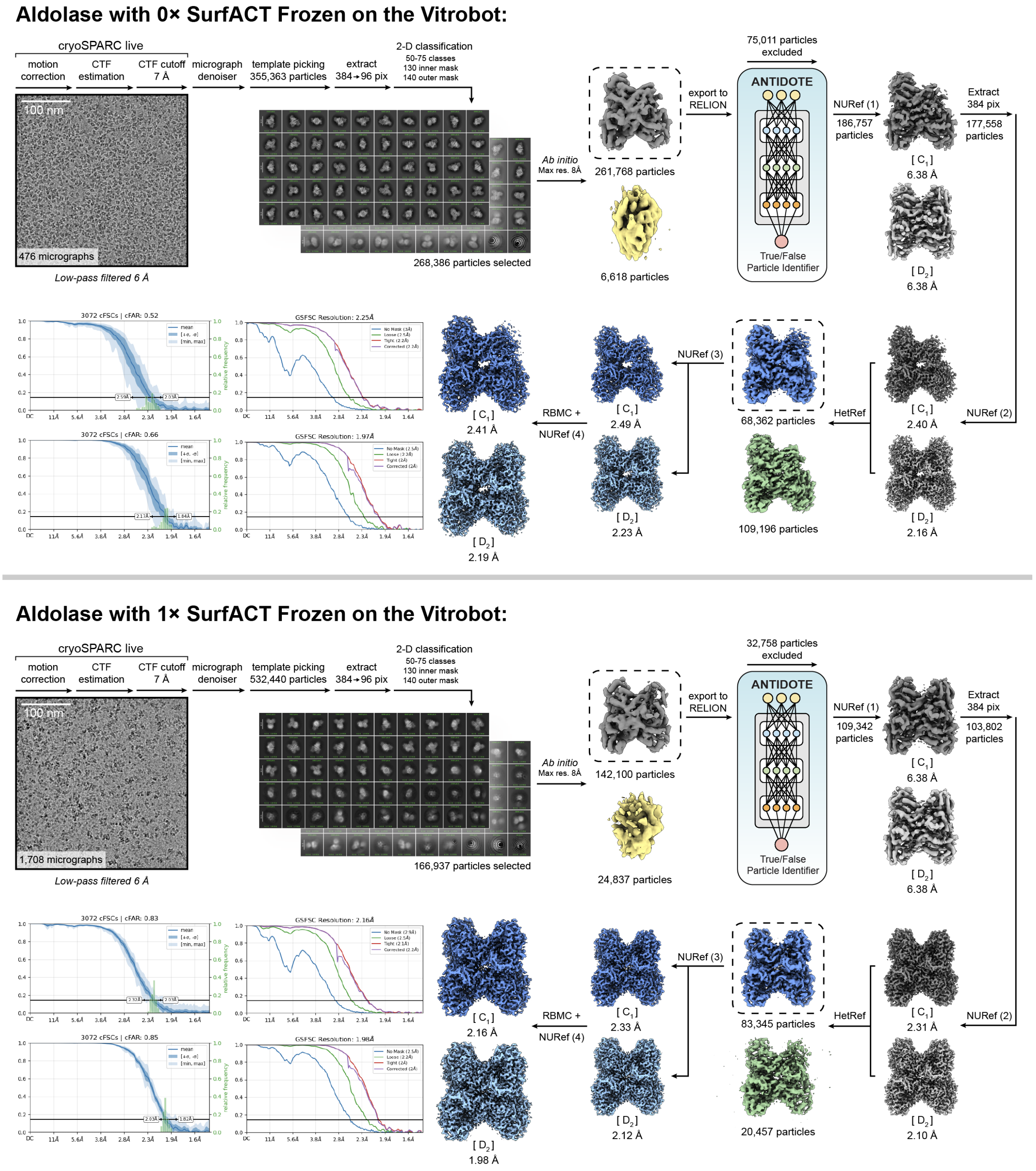


**Non-uniform refinement (NURef) 1**: minimize over per-particle scale, initialize noise model from images. **NURef 2-3**: minimize over per-particle scale, initialize noise model from images, optimize per-particle defocus, optimize CTF parameters. **NURef 4**: no real-space window, minimize over per-particle scale, initialize noise model from images, optimize per-particle defocus, optimize CTF parameters. **Heterogeneous refinement (HetRef)**: batch size per class 2000. **Reference-Based Motion Correction (RBMC):** override EER number of fractions (80).

**Figure S15: CryoEM processing of rabbit muscle aldolase prepared with 0× and 1× SurfACT using a Vitrobot Mark IV**
Single particle cryoEM workflow for aldolase with 0× and 0.1× SurfACT prepared using a Vitrobot Mark IV. Representative aligned micrographs, 2-D class averages, and 3-D sorting and refinement steps in cryoSPARC and RELION are detailed.[1,2] ANTIDOTE[3] particle curation within RELION prior to final 3-D sorting from heterogenous refinement (HetRef) in cryoSPARC with broken ($C_1$) and intact ($D_2$) volumes supplied. Final $C_1$ and $D_2$ refinements, elevation plots, and 3DFSC curves are shown. Key parameters for non-uniform refinement (NUref)[4] and reference-based motion correction (RBMC) are shown.

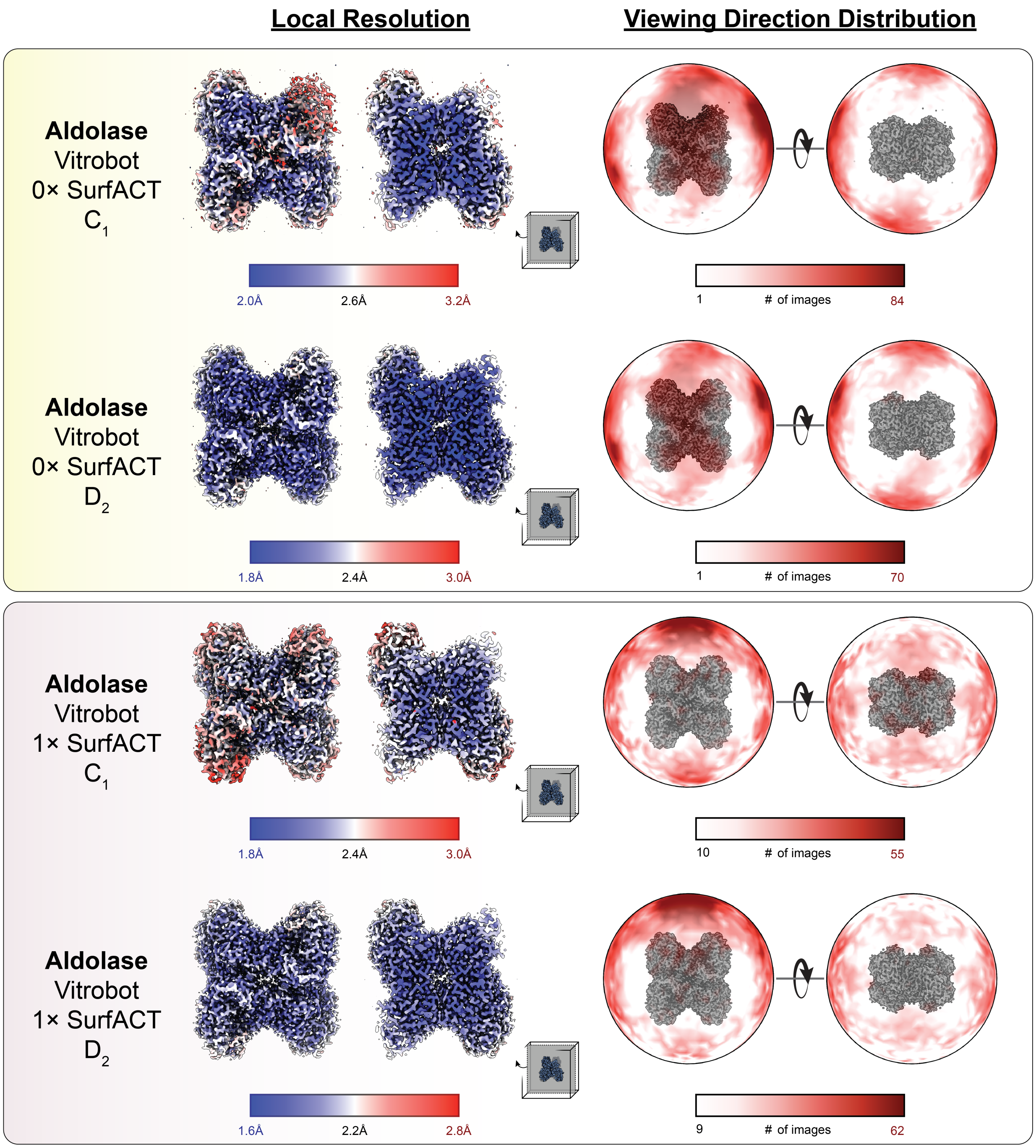


**Figure S16: Local Resolution and signal distribution for rabbit muscle aldolase prepared with 0× and 1× SurfACT using a Vitrobot Mark IV**
EM density analysis for single particle cryoEM $C_1$ and $D_2$ reconstructions of rabbit muscle aldolase with 0× and 1× SurfACT prepared using a Vitrobot Mark IV. Full volume and central slice of volume (*left*) are colored by local resolution blue (low)-white-red (high). Signal distribution (*right*) is shown as a heat map from white (low) to red (high).

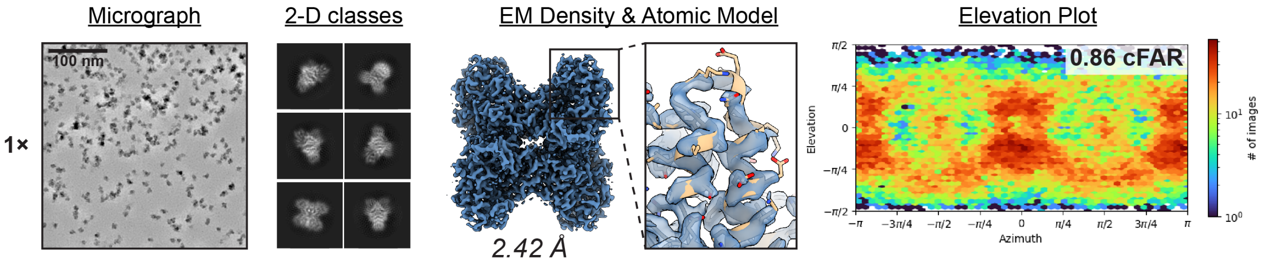


**Figure S17: Structure and properties of aldolase prepared with 1× SurfACT using a SPT Labtech chameleon with gold-coated nanowire grids**
Representative denoised micrograph of rabbit muscle aldolase frozen with 1× SurfACT using a SPT Labtech chameleon with gold-coated nanowire grids. Representative 2-D class averages, 2.4 Å aldolase EM density ($C_1$ symmetry; blue surface), flyout of model-to-density agreement of a peripheral $\alpha$-helix of destabilized protomer, and cryoSPARC[1] elevation plot to demonstrate signal distribution are shown.

### Aldolase with 1× SurfACT Frozen on the chameleon on Gold-Coated Nanowire Grids:

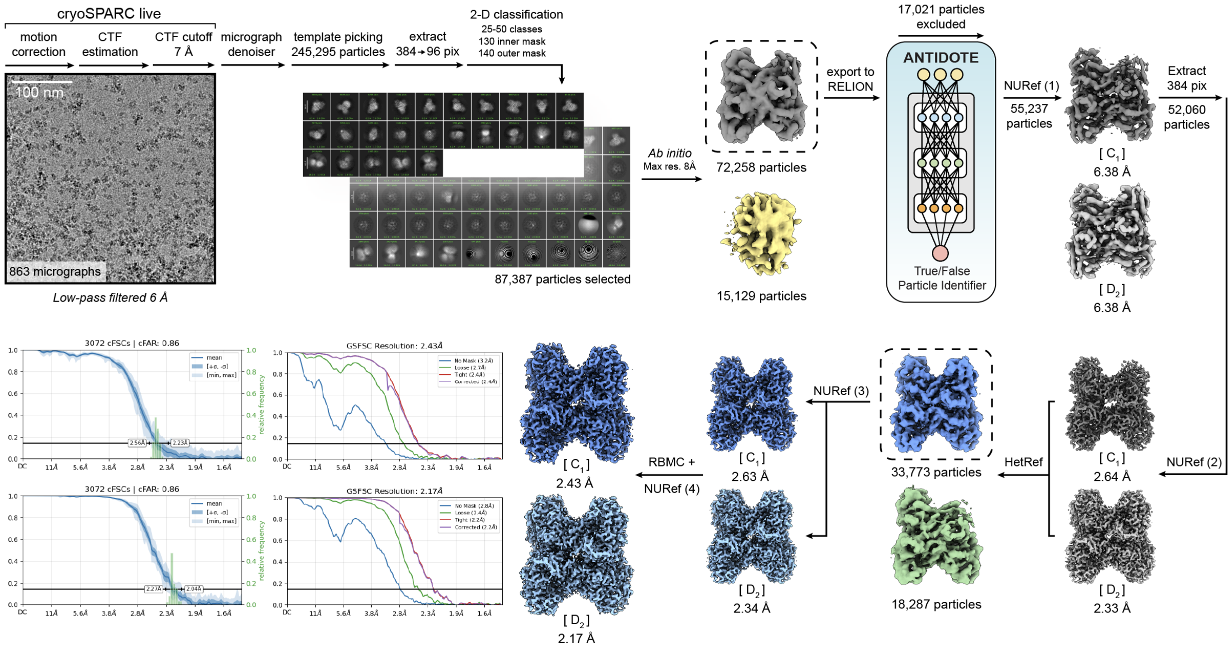


**Non-uniform refinement (NURef) 1**: minimize over per-particle scale, initialize noise model from images. **NURef 2-3**: minimize over per-particle scale, initialize noise model from images, optimize per-particle defocus, optimize CTF parameters. **NURef 4**: no real-space window, minimize over per-particle scale, initialize noise model from images, optimize per-particle defocus, optimize CTF parameters. **Heterogeneous refinement (HetRef)**: batch size per class 2000. **Reference-Based Motion Correction (RBMC):** override EER number of fractions (80).

**Figure S18: CryoEM processing of rabbit muscle aldolase prepared with 1× SurfACT using a SPT Labtech chameleon with gold-coated nanowire grids**
Single particle cryoEM workflow for aldolase with 1× SurfACT prepared using a SPT Labtech chameleon with gold-coated nanowire grids. Representative aligned micrographs, 2-D class averages, and 3-D sorting and refinement steps in cryoSPARC and RELION are detailed.[1,2] ANTIDOTE[3] particle curation within RELION prior to final 3-D sorting from heterogenous refinement (HetRef) in cryoSPARC with broken ($C_1$) and intact ($D_2$) volumes supplied. Final $C_1$ and $D_2$ refinements, elevation plots, and 3DFSC curves are shown. Key parameters for non-uniform refinement (NUref)4[4] and reference-based motion correction (RBMC) are shown.

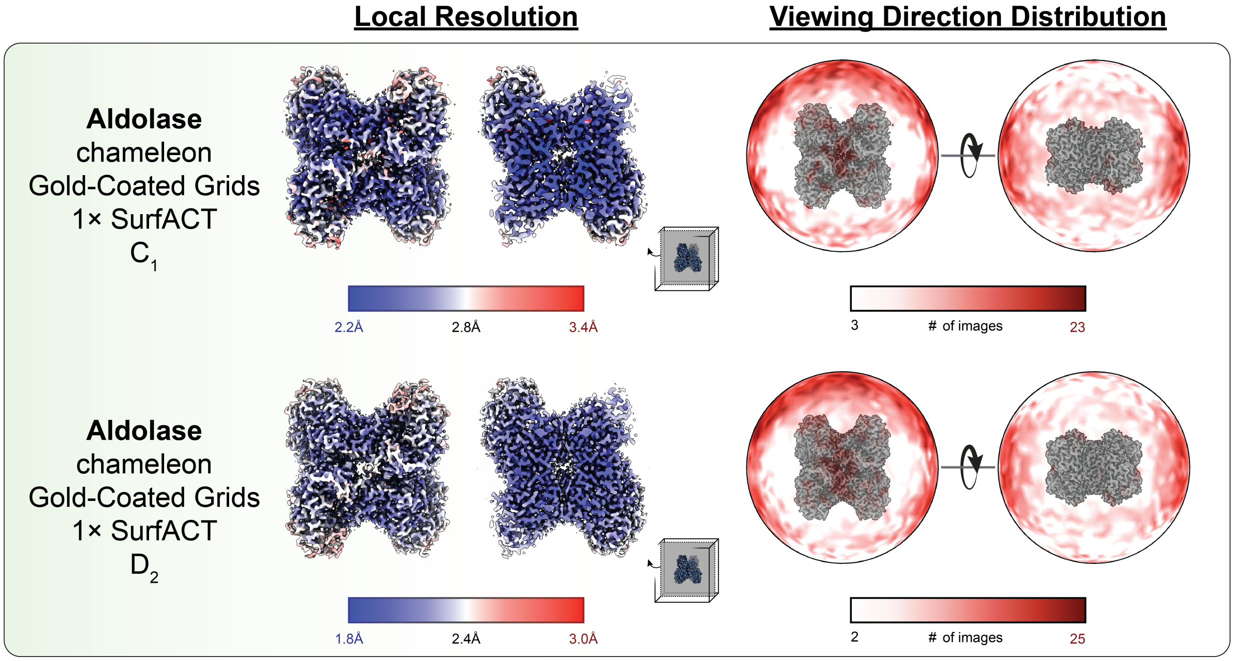


**Figure S19: Local Resolution and signal distribution for rabbit muscle aldolase prepared with 1× SurfACT using a SPT Labtech chameleon and gold-coated nanowire grids**

EM density analysis for single particle cryoEM $C_1$ and $D_2$ reconstructions of rabbit muscle aldolase with 1× SurfACT prepared using a SPT Labtech chameleon and gold-coated nanowire grids. Full volume and central slice of volume (*left*) are colored by local resolution blue (low)-white-red (high). Signal distribution (*right*) is shown as a heat map from white (low) to red (high).

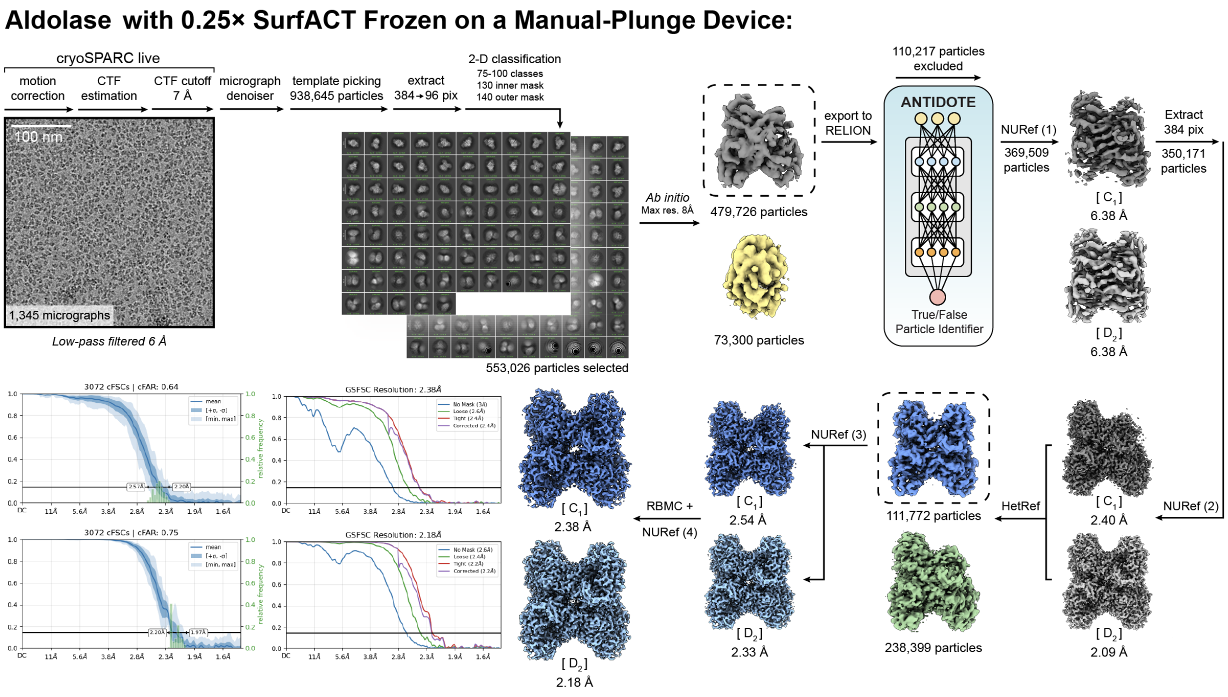


**Non-uniform refinement (NURef) 1**: minimize over per-particle scale, initialize noise model from images. **NURef 2-3**: minimize over per-particle scale, initialize noise model from images, optimize per-particle defocus, optimize CTF parameters. **NURef 4**: no real-space window, minimize over per-particle scale, initialize noise model from images, optimize per-particle defocus, optimize CTF parameters. **Heterogeneous refinement (HetRef)**: batch size per class 2000. **Reference-Based Motion Correction (RBMC):** override EER number of fractions (80).

**Figure S20: CryoEM processing of rabbit muscle aldolase prepared with 0.25× SurfACT using a manually operated plunging device**
Single particle cryoEM workflow for aldolase with 0.25× SurfACT prepared using a manually operated plunging device. Representative aligned micrographs, 2-D class averages, and 3-D sorting and refinement steps in cryoSPARC and RELION are detailed.[1,2] ANTIDOTE[3] particle curation within RELION prior to final 3-D sorting from heterogenous refinement (HetRef) in cryoSPARC with broken ($C_1$) and intact ($D_2$) volumes supplied. Final $C_1$ and $D_2$ refinements, elevation plots, and 3DFSC curves are shown. Key parameters for non-uniform refinement (NUref)[4] and reference-based motion correction (RBMC) are shown.

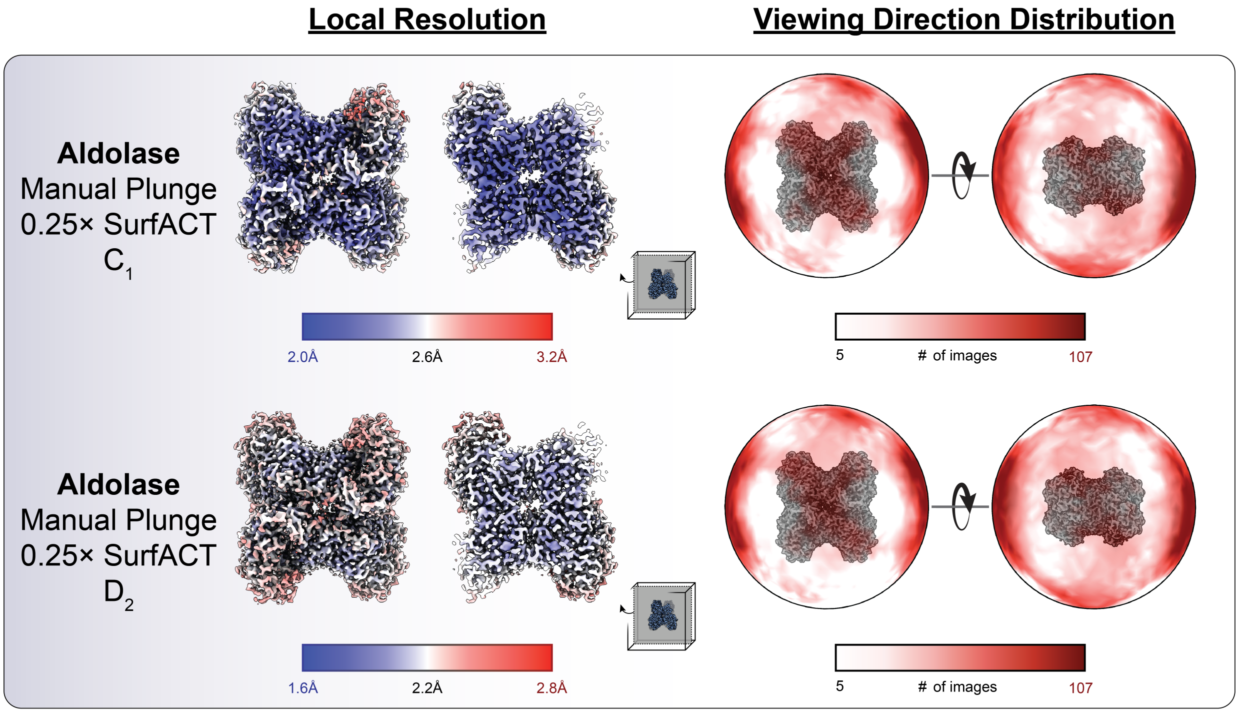


**Figure S21: Local Resolution and signal distribution for rabbit muscle aldolase prepared with 0.25× SurfACT using a manually operated plunging device**
EM density analysis for single particle cryoEM $C_1$ and $D_2$ reconstructions of rabbit muscle aldolase with 0.25× SurfACT using a manually operated plunging device. Full volume and central slice of volume (*left*) are colored by local resolution blue (low)-white-red (high). Signal distribution (*right*) is shown as a heat map from white (low) to red (high).

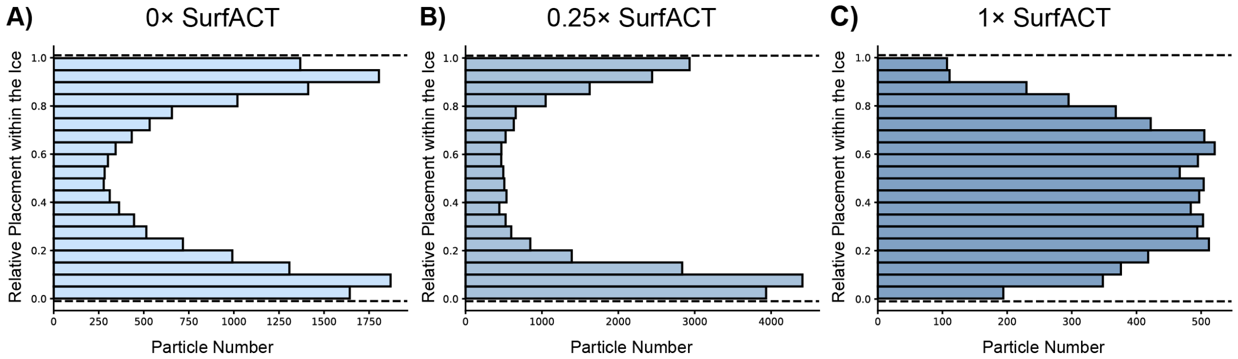


**Figure S22: Aldolase STA Particle Localization Relative to the AWI**
Quantitative analysis of particle placement relative to tomogram ice boundaries by histogram. **A**) With 0× SurfACT, particles were identified mostly near the ice boundaries with few particles identified in the center of the ice. **B**) With 0.25× SurfACT, particles were similarly present both at the AWIs and in the center ice. C) With 1× SurfACT, the vast majority of particles were distributed throughout the bulk ice and away from either AWI.

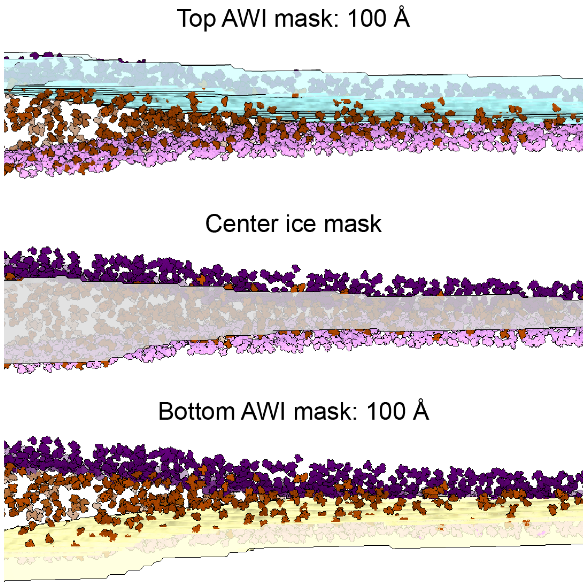


| | | tomogram concentration (mg/ml) | concentration relative to applied sample (fold) |
|---|---|---|---|
| **0x SurfACT** 6 mg/ml applied sample | **overall** | **14.0** | **2.33** |
| | top AWI | 16.4 | 2.73 |
| | center ice | 5.1 | 0.84 |
| | bottom AWI | 17.9 | 2.98 |
| | **overall AWI** | **17.1** | **2.85** |
| **0.25x SurfACT** 12 mg/ml applied sample | **overall** | **10.2** | **0.86** |
| | top AWI | 10.5 | 0.88 |
| | center ice | 3.4 | 0.28 |
| | bottom AWI | 17.3 | 1.44 |
| | **overall AWI** | **13.9** | **1.16** |
| **1x SurfACT** 12 mg/ml applied sample | **overall** | **3.6** | **0.30** |
| | top AWI | 0.7 | 0.05 |
| | center ice | 7.1 | 0.59 |
| | bottom AWI | 1.6 | 0.14 |
| | **overall AWI** | **1.2** | **0.10** |

**Figure S23: Observation that particles concentrate at the AWI that correlates with preferred orientation**
Calculation of aldolase particle concentrations in 0×, 0.25×, and 1× SurfACT tomograms. Concentrations were calculated based on volume of the mask used to define vitreous ice boundaries and the particle count contained within that mask. Overall concentration was calculated based on the complete mask. The top and bottom AWI were segmented to define 100Å from each ice surface, and areas further than 100Å from both surfaces were considered center ice (*left*). Chart shows aldolase concentration values for the overall tomogram, top AWI, center ice, bottom AWI, and both AWIs overall (*right*). Concentrations are the mean values of 5-6 tomogram datasets for each SurfACT concentration. Particle density was then compared to the applied sample concentration during grid preparation (6 mg/ml for 0× SurfACT and 12 mg/ml for 0.25× and 1× SurfACT). In 1× SurfACT tomograms, the AWIs were not marked with aldolase particles, so we could not reliably define the ice boundary for mask generation. We therefore rely on the center ice particle concentration for any conclusions, where we are confident there are no empty voxels.

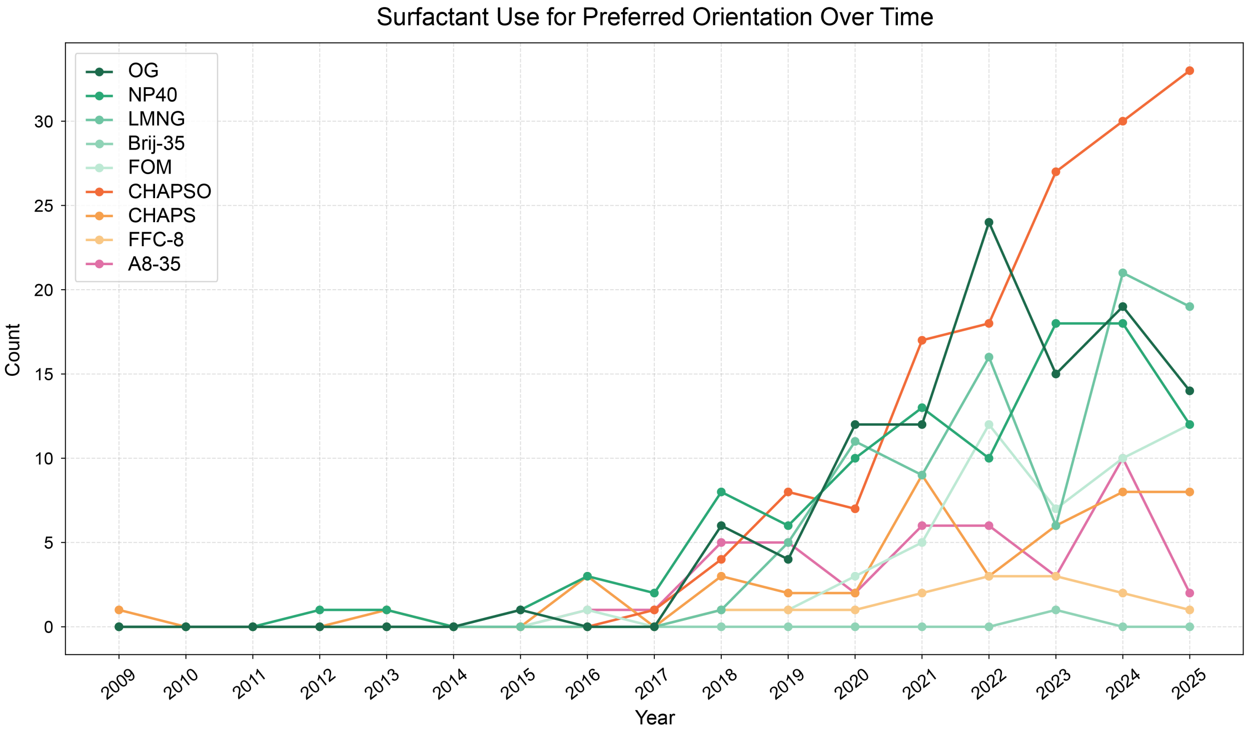


**Figure S24: Surfactant Use by Year in the EMDB**

Frequency of surfactant use for preferred orientation in EMDB-associated publications plotted by year. Surfactants include: octyl glucoside (OG), nonidet P-40 (NP40), lauryl maltose neopentyl glycol (LMNG), fluorinated octyl maltoside (FOM), Brij-35, 3-[(3-cholamidopropyl)-dimethylammonio]-2-hydroxy-1-propanesulfonate (CHAPSO), 3-[(3-cholamidopropyl)-dimethylammonio]-1-propane sulfonate (CHAPS), fluorinated fos-choline-8 (FFC-8), and amphipol 8-35 (A8-35). Plot is colored by surfactant class; non-ionic detergents are shown in shades of green, zwitterionic in shades of orange, and amphiphilic in pink.

**Table S2:** Data Collection and Processing Statistics for Hemagglutinin (A/Darwin/6/2021 H3N2)

| | Hemagglutinin (A/Darwin/6/2021 H3N2) 0× SurfACT | | Hemagglutinin (A/Darwin/6/2021 H3N2) 0.25× SurfACT | | Hemagglutinin (A/Darwin/6/2021 H3N2) 1× SurfACT | |
|---|---|---|---|---|---|---|
| **Data Collection** | | | | | | |
| Magnification | 165kx | | 165kx | | 165kx | |
| Voltage (kV) | 300 | | 300 | | 300 | |
| Spherical Aberration (mm) | 2.7 | | 2.7 | | 2.7 | |
| Electron Exposure ($e^{-}/Å^{2}$) | 65 | | 65 | | 65 | |
| Defocus range ($\mu$m; nominal) | -1 to -2.5 | | -1 to -2.5 | | -1 to -2.5 | |
| Pixel size (Å) | 0.735 | | 0.735 | | 0.735 | |
| Energy Filter Slit Width (eV) | 10 | | 10 | | 10 | |
| **Map Statistics and Post-Processing** | | | | | | |
| Particles | 94355 | 94355 | 90718 | 90718 | 35526 | 35526 |
| Symmetry imposed | $C_1$ | $C_3$ | $C_1$ | $C_3$ | $C_1$ | $C_3$ |
| Map Resolution (Å) | 2.82 | 2.65 | 2.39 | 2.16 | 2.67 | 2.47 |
| Local resolution range for 75% of voxels | 6.900 | 6.365 | 6.315 | 5.228 | 7.541 | 6.053 |
| Local resolution range (model) | 2.402-45.916 | 1.578-42.589 | 2.126-38.973 | 1.951-34.762 | 2.397-43.003 | 2.256-38.699 |
| Map sharpening B factor ($Å^{2}$) | 67.9 | 68.4 | 60.1 | 60.0 | 60.7 | 68.9 |
| Map sharpening method | Resolve | Resolve | Resolve | Resolve | Resolve | Resolve |
| Q-Score | | | 0.657 | 0.695 | 0.602 | 0.644 |
| cFAR | 0.01 | 0.01 | 0.87 | 0.9 | 0.86 | 0.9 |
| Movies | 495 | 495 | 1484 | 1484 | 1461 | 1461 |
| **Model Statistics and Validation** | | | | | | |
| Model composition | | | | | | |
| Non-hydrogen atoms | | | 13036 | 13520 | 12307 | 12654 |
| Protein | | | 1467 | 1467 | 1467 | 1467 |
| Waters | | | 1483 | 1967 | 754 | 1101 |
| R.M.S deviations | | | | | | |
| Length (Å) | | | 0.005 | 0.005 | 0.003 | 0.005 |
| Angles (°) | | | 0.836 | 0.810 | 0.599 | 0.774 |
| MolProbity score | | | 1.05 | 0.98 | 1.26 | 0.95 |
| MolProbity Clashscore | | | 2.19 | 1.97 | 3.86 | 1.84 |
| CaBLAM (% outliers) | | | 1.8 | 1.59 | 1.59 | 1.8 |
| Rotamer outliers (%) | | | 0.23 | 0.31 | 0.16 | 0.70 |
| Cis peptides (% Proline, % General) | | | 0.0/0.0 | 0.0/0.0 | 0.0/0.0 | 0.0/0.0 |
| Ramachandran Plot | | | | | | |
| Favored | | | 97.73 | 97.94 | 97.59 | 98.35 |
| Allowed | | | 2.27 | 2.06 | 2.41 | 1.65 |
| Outliers | | | 0.0 | 0.0 | 0.0 | 0.0 |

**Table S3:** Data Collection and Processing Statistics for Hemagglutinin (A/California/04/2009 H1N1)

| | Hemagglutinin (A/California/04/2009 H1N1) 0× SurfACT | | Hemagglutinin (A/California/04/2009 H1N1) 0.25× SurfACT | |
|---|---|---|---|---|
| **Data Collection** | | | | |
| Magnification | 165kx | | 165kx | |
| Voltage (kV) | 300 | | 300 | |
| Spherical Aberration (mm) | 2.7 | | 2.7 | |
| Electron Exposure ($e^-/Å^2$) | 65 | | 65 | |
| Defocus range ($\mu$m; nominal) | -1 to -2.5 | | -1 to -2.5 | |
| Pixel size (Å) | 0.735 | | 0.735 | |
| Energy Filter Slit Width (eV) | 10 | | 10 | |
| **Map Statistics and Post-Processing** | | | | |
| Particles | 24620 | 24620 | 26950 | 26950 |
| Symmetry imposed | $C_1$ | $C_3$ | $C_1$ | $C_3$ |
| Map Resolution (Å) | 3.5 | 2.84 | 2.94 | 2.64 |
| Local resolution range for 75% of voxels | 9.742 | 7.619 | 8.412 | 6.639 |
| Local resolution range (model) | 3.056-54.755 | 2.538-46.176 | 2.638-46.830 | 1.578-42.675 |
| Map sharpening B factor ($Å^2$) | 51.6 | 57.1 | 61.7 | 73.6 |
| Map sharpening method | Resolve | Resolve | Resolve | Resolve |
| Q-Score | | | 0.553 | 0.595 |
| cFAR | 0.01 | 0.04 | 0.77 | 0.82 |
| Movies | 321 | 321 | 1966 | 1966 |
| **Model Statistics and Validation** | | | | |
| Model composition | | | | |
| Non-hydrogen atoms | | | 12106 | 12401 |
| Protein | | | 1470 | 1470 |
| Waters | | | 466 | 761 |
| R.M.S deviations | | | | |
| Length (Å) | | | 0.007 | 0.005 |
| Angles (°) | | | 0.911 | 0.744 |
| MolProbity score | | | 1.34 | 1.04 |
| MolProbity Clashscore | | | 3.84 | 2.53 |
| CaBLAM (% outliers) | | | 1.66 | 1.87 |
| Rotamer outliers (%) | | | 1.09 | 1.01 |
| Cis peptides (% Proline, % General) | | | 0.0/0.0 | 0.0/0.0 |
| Ramachandran Plot | | | | |
| Favored | | | 97.26 | 98.56 |
| Allowed | | | 2.74 | 1.44 |
| Outliers | | | 0.0 | 0.0 |

**Table S4:** Data Collection and Processing Statistics for Hemagglutinin (A/Darwin/6/2021 H3N2), Subtomogram Averaged

| | Hemagglutinin (A/Darwin/6/2021 H3N2) 0× SurfACT | Hemagglutinin (A/Darwin/6/2021 H3N2) 0.25× SurfACT |
|---|---|---|
| **Data Collection** | | |
| Magnification | 64kx | 64kx |
| Voltage (kV) | 300 | 300 |
| Spherical Aberration (mm) | 2.7 | 2.7 |
| Tilt Range (Increment) | $-54°$ to $54°$, $3°$ | $-54°$ to $54°$, $3°$ |
| Total Accumulated Dose ($e^-/Å^2$) | 68 | 68 |
| Average Dose per Tilt ($e^-/Å^2$) | $\sim$1.85 | $\sim$1.85 |
| Defocus ($\mu$m; nominal) | -3.8 | -3.8 |
| Pixel size (Å) | 0.735 | 0.735 |
| Energy Filter Slit Width (eV) | 10 | 10 |
| **Map Statistics and Post-Processing** | | |
| Tomograms | 3 | 12 |
| Subtomograms | 20267 | 10359 |
| Symmetry | $C_3$ | $C_3$ |
| Resolution (Å) | 7.0 | 7.50 |

**Table S5:** Data Collection and Processing Statistics for MoFeP

| | MoFeP 0× SurfACT | | MoFeP 0.25× SurfACT | | MoFeP 1× SurfACT | |
|---|---|---|---|---|---|---|
| **Data Collection** | | | | | | |
| Magnification | 165kx | | 165kx | | 165kx | |
| Voltage (kV) | 300 | | 300 | | 300 | |
| Spherical Aberration (mm) | 2.7 | | 2.7 | | 2.7 | |
| Electron Exposure ($e^-/Å^2$) | 65 | | 65 | | 65 | |
| Defocus range ($\mu$m; nominal) | -1 to -2.5 | | -1 to -2.5 | | -1 to -2.5 | |
| Pixel size (Å) | 0.735 | | 0.735 | | 0.735 | |
| Energy Filter Slit Width (eV) | 10 | | 10 | | 10 | |
| **Map Statistics and Post-Processing** | | | | | | |
| Particles | 48236 | 48236 | 100372 | 100372 | 23561 | 23561 |
| Symmetry imposed | $C_1$ | $C_2$ | $C_1$ | $C_2$ | $C_1$ | $C_2$ |
| Map Resolution (Å) | 2.69 | 2.46 | 2.16 | 2.06 | 2.49 | 2.34 |
| Local resolution range for 75% of voxels | 7.919 | 6.890 | 6.053 | 5.394 | 7.498 | 6.640 |
| Local resolution range (model) | 2.305-43.024 | 2.208-42.869 | 1.850-35.118 | 1.773-35.030 | 2.200-42.961 | 1.589-39.013 |
| Map sharpening B factor ($Å^2$) | 52.8 | 55.3 | 49.0 | 50.1 | 49.9 | 52.3 |
| Map sharpening method | Resolve | Resolve | Resolve | Resolve | Resolve | Resolve |
| Q-Score | 0.46 | 0.558 | 0.698 | 0.733 | 0.628 | 0.669 |
| cFAR | 0.29 | 0.33 | 0.68 | 0.74 | 0.8 | 0.78 |
| Movies | 968 | 968 | 1182 | 1182 | 1080 | 1080 |
| **Model Statistics and Validation** | | | | | | |
| Model composition | | | | | | |
| Non-hydrogen atoms | 16626 | 17048 | 18465 | 18652 | 17707 | 17870 |
| Protein | 1998 | 1998 | 1998 | 1998 | 1998 | 1998 |
| Ligand1 | ICS: 2 | ICS: 2 | ICS: 2 | ICS: 2 | ICS: 2 | ICS: 2 |
| Ligand2 | CLF: 2 | CLF: 2 | CLF: 2 | CLF: 2 | CLF: 2 | CLF: 2 |
| Ligand3 | HCA: 2 | HCA: 2 | HCA: 2 | HCA: 2 | HCA: 2 | HCA: 2 |
| Ligand4 | FE: 2 | FE: 2 | FE: 2 | FE: 2 | FE: 2 | FE: 2 |
| Waters | 602 | 1027 | 2438 | 2625 | 1680 | 1843 |
| R.M.S deviations | | | | | | |
| Length (Å) | 0.009 | 0.006 | 0.006 | 0.006 | 0.006 | 0.007 |
| Angles (°) | 1.579 | 0.941 | 1.178 | 1.179 | 1.168 | 1.472 |
| MolProbity score | 1.66 | 1.66 | 1.07 | 1.09 | 1.10 | 1.16 |
| MolProbity Clashscore | 7.99 | 6.82 | 2.65 | 2.78 | 2.59 | 2.46 |
| CaBLAM (% outliers) | 1.16 | 1.66 | 0.66 | 0.45 | 0.66 | 0.66 |
| Rotamer outliers (%) | 1.10 | 0.93 | 0.58 | 0.17 | 0.64 | 0.58 |
| Cis peptides (% Proline, % General) | 4.3/0.0 | 0.0/0.0 | 6.5/0.1 | 6.5/0.1 | 6.5/0.1 | 5.4/0.1 |
| Ramachandran Plot | | | | | | |
| Favored | 96.83 | 95.88 | 97.94 | 97.89 | 97.74 | 97.29 |
| Allowed | 3.17 | 4.12 | 2.01 | 2.11 | 2.21 | 2.66 |
| Outliers | 0.0 | 0.0 | 0.05 | 0.0 | 0.05 | 0.05 |

**Table S6:** Data Collection and Processing Statistics for Aldolase

| | Aldolase Chameleon 0× SurfACT | | Aldolase Chameleon 0.25× SurfACT | | Aldolase Chameleon 0.5× SurfACT | |
|---|---|---|---|---|---|---|
| **Data Collection** | | | | | | |
| Magnification | 165kx | | 165kx | | 165kx | |
| Voltage (kV) | 300 | | 300 | | 300 | |
| Spherical Aberration (mm) | 2.7 | | 2.7 | | 2.7 | |
| Electron Exposure ($e^-/Å^2$) | 65 | | 65 | | 65 | |
| Defocus range ($\mu$m; nominal) | -1 to -2.5 | | -1 to -2.5 | | -1 to -2.5 | |
| Pixel size (Å) | 0.735 | | 0.735 | | 0.735 | |
| Energy Filter Slit Width (eV) | 10 | | 10 | | 10 | |
| **Map Statistics and Post-Processing** | | | | | | |
| Particles | 90440 | 90440 | 79923 | 77923 | 50256 | 50256 |
| Symmetry imposed | $C_1$ | $D_2$ | $C_1$ | $D_2$ | $C_1$ | $D_2$ |
| Map Resolution (Å) | 2.42 | 2.27 | 2.4 | 2.17 | 2.41 | 2.19 |
| Local resolution range for 75% of voxels | 6.69 | 5.134 | 6.574 | 5.227 | 7.072 | 5.498 |
| Local resolution range (model) | 2.166-38.877 | 1.995-34.542 | 2.144-39.035 | 1.944-34.711 | 1.589-39.073 | 1.975-34.854 |
| Map sharpening B factor ($Å^2$) | 63.0 | 70.4 | 62.4 | 63.7 | 56.5 | 59.9 |
| Map sharpening method | Resolve | Resolve | Resolve | Resolve | Resolve | Resolve |
| Q-Score | 0.629 | 0.677 | 0.637 | 0.7 | 0.623 | 0.697 |
| cFAR | 0.61 | 0.71 | 0.72 | 0.82 | 0.8 | 0.85 |
| Movies | 1137 | 1137 | 1012 | 1012 | 1039 | 1039 |
| **Model Statistics and Validation** | | | | | | |
| Model composition | | | | | | |
| Non-hydrogen atoms | 11503 | 11843 | 11352 | 11912 | 11217 | 11853 |
| Protein | 1372 | 1372 | 1372 | 1372 | 1372 | 1372 |
| Waters | 1031 | 1371 | 880 | 1440 | 745 | 1381 |
| R.M.S deviations | | | | | | |
| Length (Å) | 0.005 | 0.005 | 0.007 | 0.005 | 0.006 | 0.005 |
| Angles (°) | 0.799 | 0.758 | 0.786 | 0.760 | 0.774 | 0.770 |
| MolProbity score | 1.35 | 1.36 | 1.38 | 1.19 | 1.42 | 1.17 |
| MolProbity Clashscore | 4.23 | 3.28 | 3.18 | 1.19 | 4.37 | 1.90 |
| CaBLAM (% outliers) | 1.33 | 1.03 | 1.47 | 1.11 | 1.55 | 1.4 |
| Rotamer outliers (%) | 1.08 | 1.17 | 1.35 | 2.07 | 1.17 | 0.90 |
| Cis peptides (% Proline, % General) | 5.6/0.0 | 5.6/0.0 | 5.6/0.0 | 5.6/0.0 | 5.6/0.0 | 5.6/0.0 |
| Ramachandran Plot | | | | | | |
| Favored | 97.36 | 96.85 | 96.99 | 97.43 | 97.14 | 96.63 |
| Allowed | 2.64 | 3.15 | 3.01 | 2.57 | 2.86 | 3.37 |
| Outliers | 0.0 | 0.0 | 0.0 | 0.0 | 0.0 | 0.0 |

**Table S7:** Data Collection and Processing Statistics for Aldolase

| | Aldolase<br>Chameleon<br>1× SurfACT | | Aldolase<br>Chameleon, Gold-coated grids<br>1× SurfACT | |
|---|---|---|---|---|
| **Data Collection** | | | | |
| Magnification | 165kx | | 165kx | |
| Voltage (kV) | 300 | | 300 | |
| Spherical Aberration (mm) | 2.7 | | 2.7 | |
| Electron Exposure ($e^-/Å^2$) | 65 | | 65 | |
| Defocus range ($\mu$m; nominal) | -1 to -2.5 | | -1 to -2.5 | |
| Pixel size (Å) | 0.735 | | 0.735 | |
| Energy Filter Slit Width (eV) | 10 | | 10 | |
| **Map Statistics and Post-Processing** | | | | |
| Particles | 73219 | 73219 | 33672 | 33672 |
| Symmetry imposed | $C_1$ | $D_2$ | $C_1$ | $D_2$ |
| Map Resolution (Å) | 2.34 | 2.1 | 2.43 | 2.17 |
| Local resolution range for 75% of voxels | 6.799 | 5.255 | 7.483 | 5.879 |
| Local resolution range (model) | 2.140-39.093 | 1.910-34.557 | 2.161-39.155 | 1.953-34.974 |
| Map sharpening B factor ($Å^2$) | 57.7 | 58.6 | 51.1 | 55.3 |
| Map sharpening method | Resolve | Resolve | Resolve | Resolve |
| Q-Score | 0.655 | 0.711 | 0.646 | 0.7 |
| cFAR | 0.88 | 0.91 | 0.86 | 0.86 |
| Movies | 986 | 986 | 755 | 755 |
| **Model Statistics and Validation** | | | | |
| Model composition | | | | |
| Non-hydrogen atoms | 11589 | 12087 | 11633 | 12066 |
| Protein | 1372 | 1372 | 1372 | 1372 |
| Waters | 1117 | 1615 | 1161 | 1594 |
| R.M.S deviations | | | | |
| Length (Å) | 0.005 | 0.007 | 0.005 | 0.005 |
| Angles (°) | 0.748 | 0.830 | 0.817 | 0.808 |
| MolProbity score | 1.31 | 1.29 | 1.22 | 1.19 |
| MolProbity Clashscore | 4.42 | 3.09 | 3.28 | 2.71 |
| CaBLAM (% outliers) | 1.62 | 1.7 | 1.47 | 1.47 |
| Rotamer outliers (%) | 0.99 | 1.26 | 1.26 | 0.81 |
| Cis peptides (% Proline, % General) | 5.6/0.0 | 5.6/0.0 | 5.6/0.0 | 5.6/0.0 |
| Ramachandran Plot | | | | |
| Favored | 97.58 | 97.43 | 97.87 | 97.29 |
| Allowed | 2.42 | 2.57 | 2.13 | 2.71 |
| Outliers | 0.0 | 0.0 | 0.0 | 0.0 |

**Table S8:** Data Collection and Processing Statistics for Aldolase

| | Aldolase Vitrobot 0× SurfACT | | Aldolase Vitrobot 1× SurfACT | | Aldolase Manual Plunge 1× SurfACT | |
|---|---|---|---|---|---|---|
| **Data Collection** | | | | | | |
| Magnification | 165kx | | 165kx | | 165kx | |
| Voltage (kV) | 300 | | 300 | | 300 | |
| Spherical Aberration (mm) | 2.7 | | 2.7 | | 2.7 | |
| Electron Exposure ($e^-/Å^2$) | 65 | | 65 | | 65 | |
| Defocus range ($\mu$m; nominal) | -1 to -2.5 | | -1 to -2.5 | | -1 to -2.5 | |
| Pixel size (Å) | 0.735 | | 0.735 | | 0.735 | |
| Energy Filter Slit Width (eV) | 10 | | 10 | | 10 | |
| **Map Statistics and Post-Processing** | | | | | | |
| Particles | 68277 | 68277 | 103622 | 103622 | 111612 | 111612 |
| Symmetry imposed | $C_1$ | $D_2$ | $C_1$ | $D_2$ | $C_1$ | $D_2$ |
| Map Resolution (Å) | 2.25 | 1.97 | 2.16 | 1.98 | 2.38 | 2.18 |
| Local resolution range for 75% of voxels | 6.483 | 5.174 | 6.286 | 4.594 | 6.203 | 4.951 |
| Local resolution range (model) | 1.945-35.154 | 1.788-31.164 | 1.929-35.200 | 1.761-30.793 | 2.150-38.530 | 1.911-34.386 |
| Map sharpening B factor ($Å^2$) | 46.7 | 47.0 | 50.9 | 54.9 | 65.2 | 69.4 |
| Map sharpening method | Resolve | Resolve | Resolve | Resolve | Resolve | Resolve |
| Q-Score | 0.665 | 0.729 | 0.7 | 0.747 | 0.661 | 0.701 |
| cFAR | 0.52 | 0.66 | 0.83 | 0.85 | 0.64 | 0.75 |
| Movies | 475 | 475 | 1676 | 1676 | 1345 | 1345 |
| **Model Statistics and Validation** | | | | | | |
| Model composition | | | | | | |
| Non-hydrogen atoms | 11974 | 12229 | 12119 | 12266 | 11725 | 12021 |
| Protein | 1372 | 1372 | 1372 | 1372 | 1372 | 1372 |
| Waters | 1502 | 1757 | 1647 | 1794 | 1253 | 1549 |
| R.M.S deviations | | | | | | |
| Length (Å) | 0.005 | 0.006 | 0.005 | 0.005 | 0.005 | 0.004 |
| Angles (°) | 0.839 | 0.923 | 0.741 | 0.848 | 0.765 | 0.753 |
| MolProbity score | 1.24 | 1.18 | 0.98 | 0.95 | 1.10 | 1.13 |
| MolProbity Clashscore | 2.75 | 2.04 | 1.28 | 1.19 | 2.23 | 1.99 |
| CaBLAM (% outliers) | 1.33 | 1.11 | 1.4 | 1.4 | 1.55 | 1.18 |
| Rotamer outliers (%) | 1.17 | 1.62 | 0.81 | 0.54 | 0.90 | 0.72 |
| Cis peptides (% Proline, % General) | 5.6/0.0 | 5.6/0.0 | 5.6/0.0 | 5.6/0.0 | 5.6/0.0 | 5.6/0.0 |
| Ramachandran Plot | | | | | | |
| Favored | 97.36 | 97.8 | 97.36 | 97.43 | 97.51 | 97.07 |
| Allowed | 2.64 | 2.2 | 2.64 | 2.57 | 2.49 | 2.93 |
| Outliers | 0.0 | 0.0 | 0.0 | 0.0 | 0.0 | 0.0 |

**Table S9:** Data Collection and Processing Statistics for Aldolase, Subtomogram Averaged

| | Aldolase 0× SurfACT | Aldolase 0.25× SurfACT | Aldolase 1× SurfACT |
|---|---|---|---|
| **Data Collection** | | | |
| Magnification | 64kx | 64kx | 64kx |
| Voltage (kV) | 300 | 300 | 300 |
| Spherical Aberration (mm) | 2.7 | 2.7 | 2.7 |
| Tilt Range (Increment) | $-54°$ to $54°$, $3°$ | $-54°$ to $54°$, $3°$ | $-54°$ to $54°$, $3°$ |
| Total Accumulated Dose ($e^-$/Å$^2$) | 68 | 68 | 68 |
| Average Dose per Tilt ($e^-$/Å$^2$) | $\sim$1.85 | $\sim$1.85 | $\sim$1.85 |
| Defocus ($\mu$m; nominal) | -3.8 | -3.8 | -3.8 |
| Pixel size (Å) | 0.735 | 0.735 | 0.735 |
| Energy Filter Slit Width (eV) | 10 | 10 | 10 |
| **Map Statistics and Post-Processing** | | | |
| Tomograms | 6 | 6 | 5 |
| Subtomograms | 20089 | 22780 | 7851 |
| Symmetry | $C_1$ | $C_1$ | $C_1$ |
| Resolution (Å) | 7.36 | 8.80 | 9.01 |

**Table S10:** Data Collection and Processing Statistics for Aldolase 0.25× SurfACT, Subtomogram Averaged

| | Aldolase 0.25× SurfACT Top air-water interface | Aldolase 0.25× SurfACT Center ice | Aldolase 0.25× SurfACT Bottom air-water interface |
|---|---|---|---|
| **Data Collection** | | | |
| Magnification | 64kx | 64kx | 64kx |
| Voltage (kV) | 300 | 300 | 300 |
| Spherical Aberration (mm) | 2.7 | 2.7 | 2.7 |
| Tilt Range (Increment) | $-54°$ to $54°$, $3°$ | $-54°$ to $54°$, $3°$ | $-54°$ to $54°$, $3°$ |
| Total Accumulated Dose ($e^-/Å^2$) | 68 | 68 | 68 |
| Average Dose per Tilt ($e^-/Å^2$) | $\sim$1.85 | $\sim$1.85 | $\sim$1.85 |
| Defocus ($\mu$m; nominal) | -3.8 | -3.8 | -3.8 |
| Pixel size (Å) | 0.735 | 0.735 | 0.735 |
| Energy Filter Slit Width (eV) | 10 | 10 | 10 |
| **Map Statistics and Post-Processing** | | | |
| Tomograms | 6 | 6 | 6 |
| Subtomograms | 4854 | 3620 | 7949 |
| Symmetry | $C_1$ | $C_1$ | $C_1$ |
| Resolution (Å) | 10.11 | 10.08 | 9.98 |
